# A Type III secretion system effector evolved to be mechanically labile and initiate unfolding from the N-terminus

**DOI:** 10.64898/2025.12.03.691928

**Authors:** Katherine E. DaPron, Alexandre M. Plastow, Morgan R. Fink, Brandon Dzuba, Marc-Andre LeBlanc, Thomas T. Perkins, Emad Tajkhorshid, Marcelo C. Sousa

## Abstract

Many Gram-negative pathogens critically depend on the Type III secretion system (T3SS) to inject effector proteins into host cells for colonization. Because the channel of the T3SS is narrow (∼2 nm), effectors must be unfolded for secretion. However, the T3SS cannot unfold mechanically robust substrates (GFP, ubiquitin, and dihydrofolate reductase), severely impairing their secretion. Consistent with this, effectors are exceptionally mechanically labile, unfolding at low forces. Thus, secretion competency is correlated with mechanical properties. Effector sequences have significantly diverged from non-effectors, suggesting that secretion exerts evolutionary pressure selecting mechanical lability. Here, using atomic-force-microscopy–based force spectroscopy, we show that effector NleC is mechanically labile (*F*_unfold_ = 13.5 pN at 100 nm/s) and mechanically compliant, as characterized by a large distance to the transition state (Δ*x*^‡^ = 2.7 nm). In contrast, the non-effector homolog protealysin is mechanically stable (*F*_unfold_ = 50.7 pN at 100 nm/s) and brittle (Δ*x*^‡^ = 0.7 nm), comparable to proteins known to impair secretion (*F*_unfold_ > 80 pN; Δ*x*^‡^ < 0.4 nm). Denaturant-induced unfolding assays demonstrate that effectors exhibit rates typical of their fold, further reinforcing mechanical properties rather than fast unfolding kinetics (*k*_0_) predicts secretion. Steered molecular dynamic simulations revealed NleC unfolding initiates at the N-terminus, consistent with current secretion models, whereas protealysin unfolding initiates at the C-terminus. Notably, the NleC N-terminus is primarily α-helical while non-effector homologs contain β-sheets, which may account for the distinct unfolding pathway. Together, these results support the notion that mechanical lability is an evolved, structurally encoded feature underlying effector secretion.

**Significance:** The Type III secretion system (T3SS) delivers effector proteins directly into host cells to promote bacterial colonization. Effectors must be unfolded for secretion, and this particular selective pressure is hypothesized to have driven significant sequence divergence from non-effector proteins. Here, we show that effectors are not characterized by unusually fast unfolding rates. Rather as hypothesized, effector NleC is more mechanically labile than its non-effector homolog, indicating that mechanical lability underlies both effector sequence divergence and T3SS unfolding. Simulations revealed that NleC unfolding initiates via the N-terminus consistent with the current secretion mechanism, while protealysin unfolds from the C-terminus. Together, these results strongly suggest mechanical lability is an evolved property of effectors and provide structural insight into how it is encoded.

## Introduction

The Type III secretion system (T3SS) is a specialized nanomachine of Gram-negative bacteria that delivers effector proteins directly into the host to facilitate colonization in symbiotic or pathogenic context. The T3SS complex resembles a nanoscale syringe that spans both the inner and outer bacterial and host cell membranes, effectively connecting their cytosols (Fig. 1 *A*). Effectors are synthesized in the bacterial cytosol. They are characterized by an unstructured N-terminal T3SS targeting sequence, followed by a loosely structured chaperone binding domain for chaperone-dependent effectors and well folded C-terminal effector domains (1-6). Effectors, alone or in their chaperone complexes, are trafficked to the base T3SS where their N-termini engage with the secretion apparatus, and the T3SS ATPase catalyzes chaperone release (7). Because the lumen of the T3SS needle is only ∼2 nm, secretion requires unfolding of effector domains. Current models propose that an effector’s N-terminus is threaded into the T3SS with subsequent unfolding and secretion powered by the ATPase and proton motive force (8, 9). Effectors refold once translocated into the host cell and thereby modulate processes, such as downplaying the immune response (10-14) or inducing dramatic cytoskeleton rearrangements (15-17).

**Figure 1:**
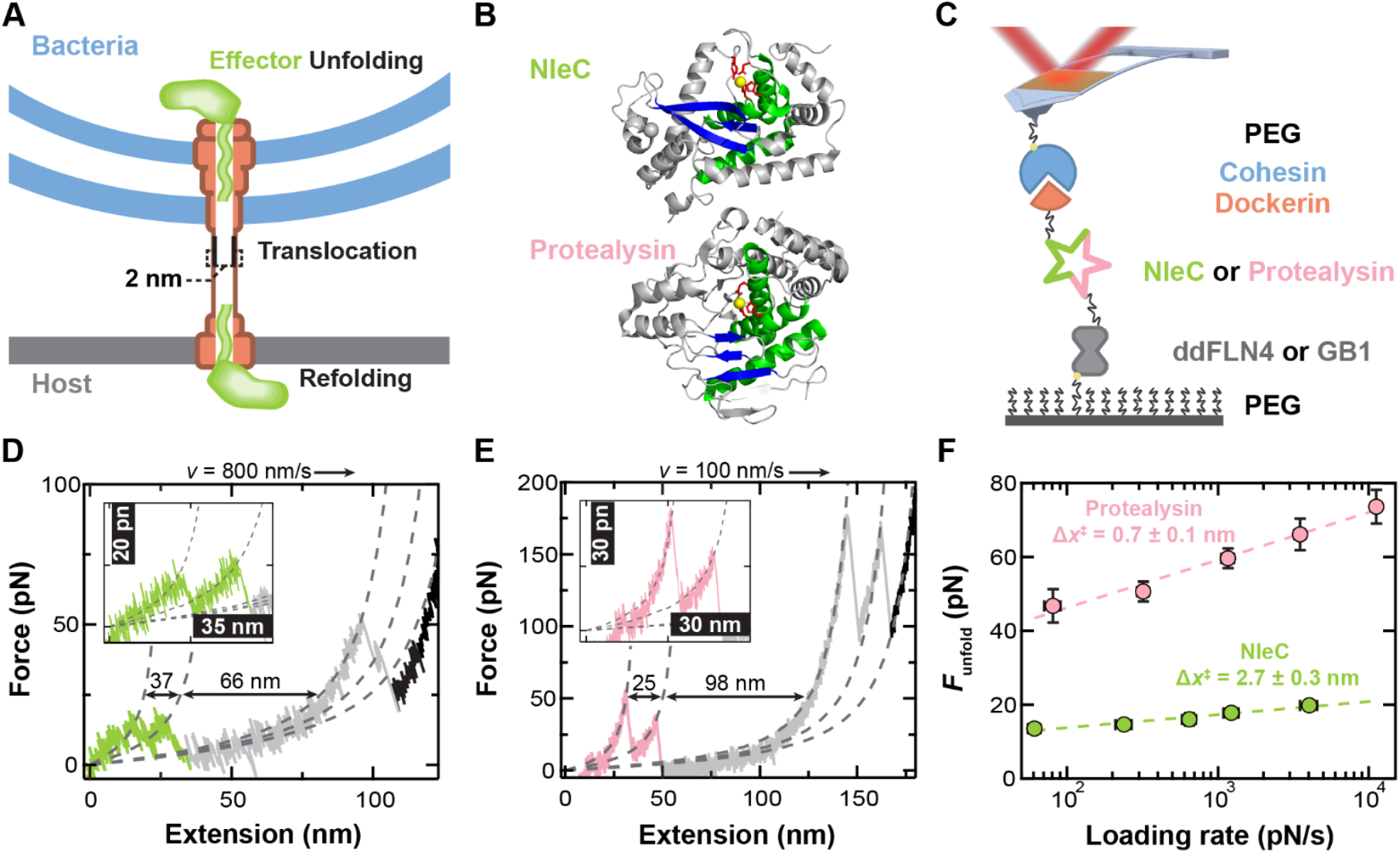
Mechanical stability of the T3SS effector NleC and non-effector homolog protealysin probed by a single-molecule atomic-force-microscopy (AFM) assay. (A) A schematic of Type III secretion. Effectors are first synthesized and folded in the bacterial cytosol. A T3SS unfoldase catalyzes effector unfolding, so they can translocate through the narrow T3SS needle (∼2 nm). Effectors then refold in the host cytosol to modulate processes. (B) Crystal structures (grey) of NleC (top; PDB 4Q3J) and protealysin (bottom; PDB 2VQX) display a highly conserved Zincin fold of a three-helix bundle (green) and a mixed β-sheet (blue) centered around a zinc-binding motif, HExxH (shown in red sticks), and catalytic zinc (yellow). (C) Cartoon showing the single-molecule assay where retraction of a focused-ion-beam modified AFM cantilever stretches the polyprotein that is site specifically anchored to a PEG-coated surface. The polyprotein encodes a well-defined marker protein (ddFLN4 or GB1, grey) followed by either the effector NleC (green) or non-effector homolog protealysin (pink), and dockerin (orange). Dockerin forms a strong, reversible protein-protein interaction with cohesin (blue) which enables stretching by a cohesin-coated cantilever. (D,E) Representative force extension curves (FECs) at constant pulling velocity depict NleC (D, green) and protealysin (E, pink) unfolding as denoted by a sharp reduction in force. Segments of the FEC were fit with a worm-like chain model (dashed lines) with the resulting change in contour length (ΔL_0_) indicated by black arrows. Data smoothed to 5 kHz. (D) FEC was collected at 800 nm/s pulling velocity and depicts NleC unfolding (green) through a single intermediate followed by the characteristic two-step unfolding of ddFLN4 (grey). (E) FEC was collected at 100 nm/s pulling velocity and shows protealysin unfolding (pink) through a single intermediate and the unfolding of two copies of GB1 (grey). (F) The dynamic force spectra of NleC (green) and protealysin (pink) shows the average unfolding force versus the logarithm of the loading rate (∂F/∂t). Data point averages were determined from at least 20 unfolding events, and their error bars represent the SEM. Fitting the Bell-Evans model (dashed line) yielded the distance to the transition state, Δx^‡^.

Effector unfolding is critical for secretion. In most cellular contexts, protein unfolding is catalyzed by AAA+ ATPases. These hexameric unfoldases can even unfold mechanically stable proteins (*e.g*., GFP and ubiquitin) (18-21). However, the T3SS ATPase does not belong to the AAA+ family. Rather, it is homologous to the *β* subunit of the F_1_ ATP synthase that operates as a rotary motor (22, 23). Therefore, the mechanism of effector unfolding remains poorly understood. It is nevertheless clear that, in contrast to AAA+ ATPases, the T3SS unfoldase cannot unfold certain substrates. Specifically, studies of effector fusions to GFP, ubiquitin, or dihydrofolate reductase (DHFR) show that secretion is strongly impaired as the fused domain resists unfolding (7, 24, 25). Structural analyses of these stalled complexes reveal that effectors are unfolded within the T3SS needle whereas the folded fusion remains trapped in the basal body of the T3SS (26, 27). These findings establish that effector unfolding is a kinetic barrier that can be rate limiting due to the limited unfolding capacity of the T3SS unfoldase compared to canonical unfoldases (*e.g*., ClpX). It has thus been proposed that effectors are thermodynamically unstable to accommodate their secretion (7, 24, 25, 28). However, recent work characterizing Salmonella effectors SopE2 and SptP showed that they have unremarkable thermodynamic stabilities, typical of globular proteins. Rather, they are instead mechanically labile (29). This result has led to the new model that mechanical stability is a determinant for secretion as effectors are mechanically labile while substrates that are known to impair secretion (GFP, ubiquitin and DHFR) are mechanically robust (29).

The requirement for effector proteins to be mechanically labile may be the origin of their extreme sequence divergence from non-effector homologs. Indeed, it typically requires three-dimensional structure analyses to identify non-effector homologs given their <15% sequence identity. Here, we directly test the hypothesis that effector proteins unfold at lower force than their non-effector homologs by comparing the mechanical stabilities of the T3SS effector NleC and its non-effector homolog protealysin. Both proteins are members of the MA clan of zinc-dependent proteases, sharing a conserved zincin fold. Using atomic force microscopy (AFM)–based single-molecule force spectroscopy (SMFS), we show that NleC unfolds at low force and is therefore mechanically labile while protealysin is mechanically robust. Accompanying steered molecular dynamics (SMD) simulations provided molecular insight by detailing unfolding pathways and demonstrated that NleC unfolds from the N-terminus. Conversely, protealysin unfolds from the C-terminus indicating that its N-terminus is mechanically robust. Finally, analysis of the NleC structure shows a divergent N-terminus in comparison with protealysin and other zincins consistent with the evolution of a mechanical labile structure to facilitate its unfolding by the T3SS unfoldase.

## Results and Discussion

### An Effector and its Non-Effector Homolog: NleC and Protealysin

To test our hypothesis that effectors may have evolved to be mechanically labile under selective pressure imposed by secretion, it was necessary to identify model effector and non-effector homolog proteins. Effectors that were previously mechanically characterized, SopE2 or SptP, do not have obvious non-effector homologs. The enterohemorrhagic *Escherichia coli* effector NleC, when delivered to the host cell, proteolytically cleaves transcription factor NF-*κ*B, thereby depressing transcription events mediating host immune responses (12-14, 30). NleC [332 amino acids (aa)] has an unstructured N-terminus (21 aa) that is rich in serine and proline residues, typical of the T3SS targeting signal of chaperone-independent effectors (1, 5, 6, 31, 32). NleC shares less than 15% sequence identity with any non-effector protein, further emphasizing the extreme sequence divergence of effectors. However, the NleC crystal structure reveals that it belongs to the MA clan of metallopeptidases [MEROPS database (33)], with a basic Zincin fold of a three-helix bundle and mixed β-sheet around a conserved zinc-binding HExxH motif (31, 34, 35) (Fig. 1 *B*). Thus, another member of the MA metallopeptidase clan would be an appropriate reference to compare mechanical properties with NleC.

Thermolysin is perhaps the best characterized member of the MA metallopeptidases. However, its larger size [548 amino acids (aa)] complicates comparisons to NleC. Nevertheless, protealysin was identified as a 341 aa thermolysin-like protease from *Serratia proteamaculans*, which does not have a T3SS. Thus, protealysin is a suitable non-effector homolog of NleC. Like NleC, the structure of protealysin has been experimentally determined, revealing a well-structured zincin fold (6–341 aa) although a short segment (residues 38–51) was not modeled due to conformational flexibility (36) (Fig. 1 *B*).

### Effector NleC is Less Mechanically Stable than its Non-Effector Homolog Protealysin

We sought to compare the mechanical properties of NleC and protealysin using a previously developed AFM-based SMFS assay (29). In this assay, a polyprotein is stretched between a glass coverslip and an AFM cantilever (Fig. 1 *C*). The polyprotein incorporates a N-terminal ybbr-tag that is covalently attached to a PEG-CoA functionalized surface via the enzymatic activity of sfp (37). Following the ybbr-tag is a ddFLN4 marker domain, which unfolds with a characteristic two-step pattern used to verify single-molecule attachment (38). Cohesin conjugated to PEG-coated cantilevers allows for site-specific attachment to the polyprotein via its mechanically robust interaction with dockerin (>300 pN) (39, 40). Site-specific anchoring of the polyprotein to both a PEG-functionalized surface and cantilever minimized non-specific attachment and adhesion, while the mechanically robust yet reversible tip attachment allowed entire datasets to be collected with a single cantilever for improved precision (41). Given the low unfolding force of effectors, we enhanced the force precision and time resolution by using focused-ion-beam-modified cantilevers (42-44).

Our single-molecule assay was initiated by gently pressing (75–100 pN) the cohesin-coated cantilever into the polyprotein-coated surface for 0.15 s. The cantilever then retracted at a constant velocity (*v*) such that the polyprotein was stretched. Protein domain unfolding resulted in a rapid drop in force (*F*). Force was detected by cantilever deflection. The resulting force-extension curves (FECs) had a characteristic sawtooth pattern, where each peak corresponded to individual unfolding events (Fig. 1 *D*). For NleC, ∼4% of cantilever retractions were consistent with stretching a single polyprotein at *v* = 100, 400, 800, 1600 and 3200 nm/s, showing NleC unfolding followed by the characteristic double-peak structure of ddFLN4. (gray in Fig. 1 *D*). While Fig. 1 *D* shows a single unfolding intermediate for NleC, its unfolding pathway was heterogenous. These FECs showed 0 (8%), 1 (42%), 2 (37%) and 3 or more (13%) intermediates at 100–3200 nm/s (*SI Appendix*, Fig. S1). To analyze these FECs, individual segments were fit to a worm-like chain (WLC) model (45) (Fig. 1 *D*, dashed lines) that models the entropic stretching of an unstructured polypeptide with a persistence length (*p* = 0.4 nm (46)) and contour length (*L*_*0*_). For NleC unfolding, the Δ*L*_*0*_ between the first peak in the FEC and the ddFLN4 marker domain was 91.6 ± 0.5 nm (mean ± SEM; *N* = 283) in good agreement with the expected value of 90.4 nm based on its structure and a *L*_*0*_ per aa of 0.365 nm/aa (47, 48). NleC unfolded at low force [13.5 ± 0.5 pN (mean ± SEM; N = 64 at 100 nm/s)]. This low unfolding force is consistent with the previously reported mechanical stabilities of effectors SopE2 and SptP [16.5 ± 1.8 pN (mean ± SEM; *N* = 31) and 12.6 ± 0.7 pN (*N* = 25) at 100 nm/s, respectively], indicating that T3SS effectors are mechanically labile (29).

In pilot experiments, protealysin showed significantly higher unfolding forces than NleC that were close in magnitude to the unfolding forces of the ddFLN4 marker domain. To facilitate analysis, we modified the polyprotein by substituting ddFLN4 for a more mechanically robust marker protein, GB1 (49). With this new AFM construct, the resulting FEC yielded easily assignable peaks with protealysin unfolding before GB1 (Fig. 1 *E*) with ∼7% of cantilever retractions yielding single-molecule attachments at *v* = 30, 100, 300, 1000 and 3000 nm/s. WLC fits showed that the Δ*L*_*0*_ between the first unfolding peak and the first GB1 domain unfolding was 126.3 ± 0.7 nm (mean ± SEM; *N* = 156) in good agreement with the expected value of 120.9 nm for protealysin unfolding. As with NleC, the FECs for protealysin frequently displayed a variable number of intermediate unfolding peaks (pink in Fig. 1 *E*), composed of 0 (18%), 1 (48%), 2 (25%), 3 or more (9%) intermediates at 30–3000 nm/s (*SI Appendix*, Fig. S2).

To compare the mechanical properties of NleC and protealysin, we measured their initial unfolding force over a range of velocities (30–3200 nm/s), yielding a dynamic force spectrum (50) (Fig. 1 *F*). In agreement with our hypothesis that effectors are mechanically labile compared to non-effector homologs, the average NleC unfolding force was more than 3-fold lower than protealysin over all velocities studied. More quantitatively, NleC unfolded at 13.5 ± 0.5 pN (mean ± SEM; *N* = 64 at 100 nm/s) while protealysin unfolded at 50.7 ± 2.7 pN (mean ± SEM; *N* = 37 at 100 nm/s), a highly significant difference (*p* < 0.001; unpaired T-test).

### Effector NleC is Mechanically Compliant

Not only were the magnitudes of the unfolding forces between NleC and protealysin distinctly different, the slopes of the two dynamic force spectrums were as well (Fig. 1 *F*). This difference in slope for the average unfolding force plotted as a function of the logarithm of loading rate (*r* = ∂*F/*∂*t*) arises from a change in the distance to the transition state (Δ*x*^‡^), as described by Bell-Evans model (51). Within SMFS, Δ*x*^‡^ can be interpreted as how much a protein deforms along the mechanical stretching reaction coordinate before unfolding. Thus, a large Δ*x*^‡^ is associated with a compliant protein, whereas a protein with small Δ*x*^‡^ is described as brittle. Of particular note for this study, GFP, ubiquitin and DHFR—all proteins that cannot be secreted by the T3SS—are brittle, with Δ*x*^‡^ of 0.28, 0.23 and 0.37 nm, respectively (47, 52, 53).

Using the Bell-Evans model, we analyzed the dynamic force spectrum for NleC and protealysin, which yielded Δ*x*^‡^ along with *k*_0_, the zero-force unfolding rate (*SI Appendix*, Table S1). For NleC, Δ*x*^‡^ was 2.7 ± 0.3 nm (mean ± fit error), representing one the most compliant proteins measured to date by AFM (54). This result is consistent with the previously characterized effectors SopE2 and SptP that also proved to be highly mechanically compliant [Δ*x*^‡^ = 1.6 ± 0.4 nm (mean ± fit error) and 1.4 ± 0.2 nm, respectively] (29). In contrast, analysis of the protealysin data yielded a Δ*x*^‡^ = 0.7 ± 0.1 nm (mean ± fit error), which is significantly closer to proteins that inhibit secretion than NleC.

Compliance is relevant to effector secretion because there is a hyperbolic relationship between Δ*x*^‡^ and unfolding forces for a set of mechanically characterized protein of diverse folds (54) (*SI Appendix*, Fig. S3). Previously, we showed that this hyperbolic relationship is described by the Bell-Evans model:

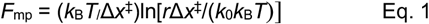

where *F*_MP_ is the most probable unfolding force, *r* is the loading rate and *k*_B_*T* is the thermal energy. This equation provides rationale for why a large Δ*x*^‡^ leads to low force unfolding (29) and underscores the central role of Δ*x*^‡^ in governing protein mechanical stability. The comparison with other proteins also highlights the selective pressure of secretion as NleC has unusually low unfolding force and high compliance for an αβ fold whereas the mechanical properties of protealysin are more representative of its fold (*SI Appendix*, Fig. S3).

The large mechanical compliance of effectors bears physiological relevance. The Bell model (55) defines unfolding rate under force (*F*) as:

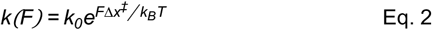

Therefore, for any given force, the unfolding rate exponentially depends on Δ*x*^‡^. The T3SS unfoldase is relatively weak and unable to unfold and secrete GFP, ubiquitin or DHFR. Nevertheless, even if small forces can be applied by the unfoldase, compliant proteins (large Δ*x*^‡^), such as those in the effectors, will unfold faster than brittle proteins (short Δ*x*^‡^). As hypothesized, NleC is highly compliant while its non-effector homolog protealysin approaches the brittleness of GFP, ubiquitin and DHFR. This result is consistent with NleC having evolved under pressure to be mechanically labile to facilitate efficient secretion by the T3SS.

### Ensemble Unfolding Kinetics of SopE2, SptP and NleC Effector Domains

While the unfolding rate under force is exponentially dependent on Δ*x*^‡^, it is also linearly dependent *k*_0_ (Eq. 2). Therefore, it is possible that effectors may have evolved fast intrinsic unfolding kinetics, rather than mechanical lability, to facilitate their secretion. It was previously reported that the *k*_0_ of SopE2 and SptP, as estimated from AFM data by fitting the Bell-Evans model to dynamic force spectra, did not correlate with secretion competency. However, these estimates require extrapolation over orders of magnitude in loading rate, which is a known problem (56). Therefore, we implemented an alternative approach. To do so, we measured the unfolding kinetics of the SopE2, SptP and NleC effector domains by circular dichroism at increasing concentrations of denaturant. The natural logarithm of the observed exponential unfolding rate constants were then plotted versus the denaturant concentration and linearly fitted to extrapolate the unfolding rate constants in the absence of denaturant (Fig. 2 *A*–*C*). This yielded *k*_0_ of 13.2 ± 1.1 x 10^-1^ (mean ± SD), 55.2 ± 1.1 x 10^-3^ and 28.7 ± 1.1 x 10^-5^ s^-1^ for SopE2, SptP and NleC, respectively.

**Figure 2:**
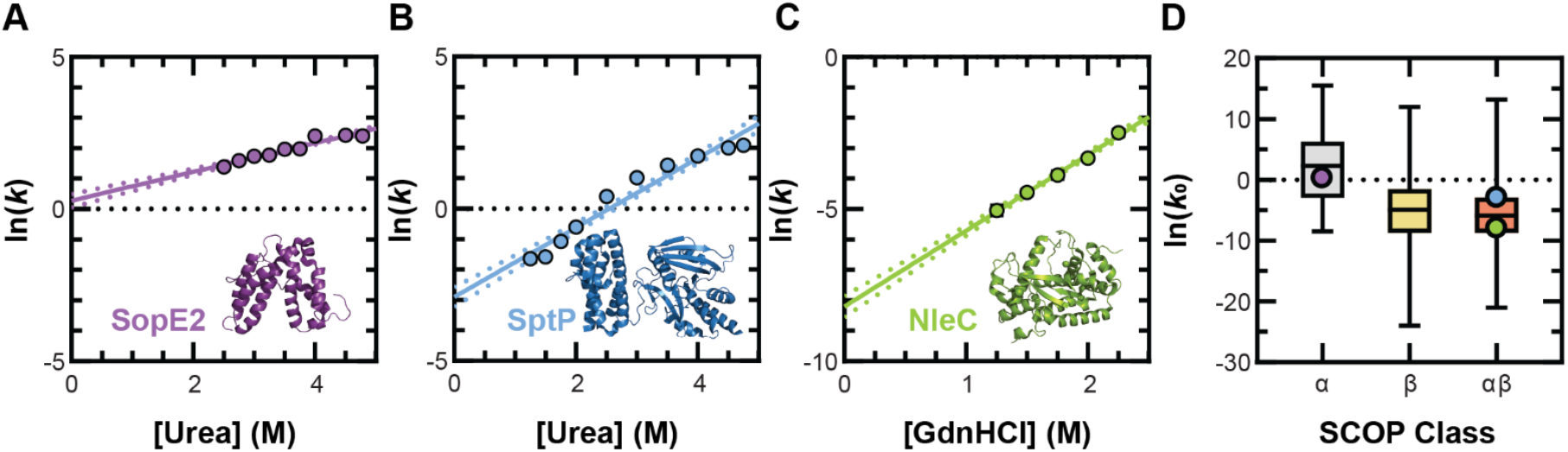
Kinetic unfolding of T3SS effector proteins. (A–C) The natural logarithm of SopE2 (A; purple), SptP (B, blue) and NleC (C, green) denaturant-dependent unfolding rates vary linearly with denaturant concentration, in which extrapolation to the absence of denaturant yields the natural logarithm of k_0_. For A–C, the average of three replicates was plotted. The error bars represent SD but are smaller than the marker size. The data was fitted with Eq. 3. Dotted lines represent the 95% confidence interval (D) Comparison of the natural logarithm of k_0_ of T3SS effectors to reference proteins of various folds. k_0_ boxplots are generated from N = 31, 37 and 40 (left to right) proteins categorized by SCOP class from reference (55). Boxes extend from 25^th^ to 75^th^ percentiles, and whiskers extend to minimum and maximum values. SopE2 (purple, all α-helical), SptP (blue, αβ), and NleC (green, αβ) are shown as individual values, representing the natural logarithm of their average k_0_. Error bars represent SD, but they are smaller than the marker size.

We then compared the observed *k*_0_ of effectors to a set of 108 proteins, representing *k*_0_ values across all α-helical, all β-sheet or αβ structural classes (57, 58). As shown in Fig. 2 *D*, the *k*_0_ of SopE2 (α-helical), SptP (αβ) and NleC (αβ) are not significantly different from those of reference proteins in the same structural class. We thus conclude that effectors do not have unusually fast intrinsic unfolding kinetics that may explain their secretion competency.

### Steered Molecular Dynamics Simulations Reveal the NleC and Protealysin Unfolding Pathways Under Force

The molecular events associated with protein unfolding in SMFS experiments can be interpreted with all-atom steered molecular dynamics (SMD) simulations (39, 59-63). In this approach, the protein structure (here atomic model) is anchored at one end while the other end is pulled using a harmonic restraint moving at constant velocity. These simulations produce trajectories that associate specific unfolding transitions with the unfolding forces (59), providing insight into the secondary and tertiary structural features responsible for mechanical stability.

We carried out three all-atom SMD simulation replicas of NleC (22–280 aa). To reproduce the pulling geometry of the AFM experiments, the N-terminal residue was harmonically restrained to its initial position, and the C-terminal residue was pulled at ∼2 Å/ns. The applied force was monitored and plotted versus the distance from the initial position to generate a simulated FEC (Fig. 3 *A*).

**Figure 3:**
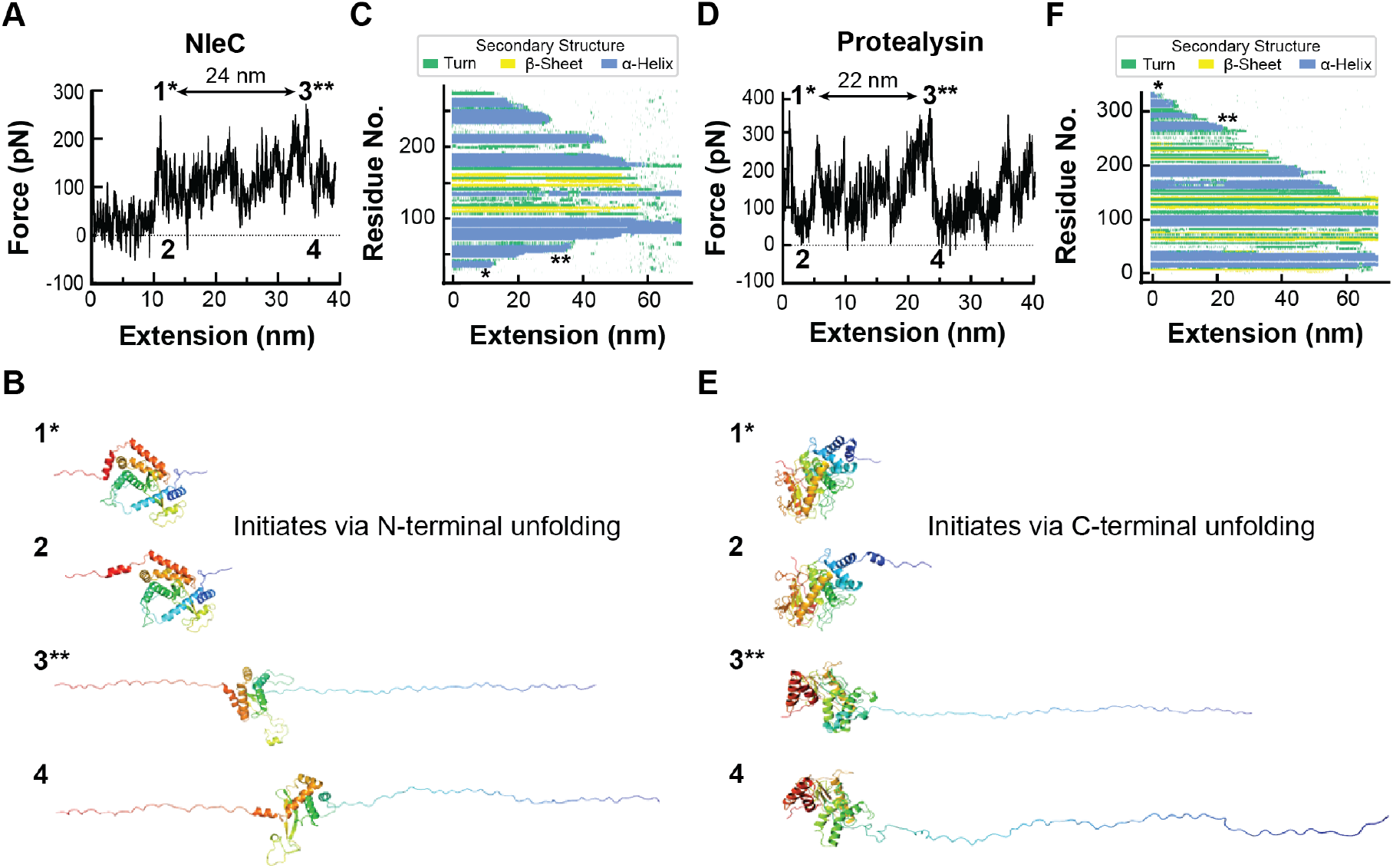
Force-induced unfolding behavior of NleC and protealysin in SMD simulations. (A and D) Representative simulated FECs for NleC (A) and protealysin (D) showing the first 40 nm of extension. The major force peaks (1* and 3**) and force drops (2 and 4) are indicated. FEC was smoothed with the Savitzky–Golay filter for visual presentation. (B and E) Cartoon representations of the atomic models in the SMD simulation frames of NleC (B) and protealysin (E) unfolding corresponding to events 1-4 indicated in (A) and (D). The proteins are colored in a spectrum from red (N-terminus) to orange to yellow to green to cyan to blue (C-terminus) to indicate directionality. (C and F) Secondary structure evolution plots of the SMD unfolding of NleC (C) and protealysin (F), respectively. Secondary structure elements in each simulation frame were colored as described in the key. FEC peaks 1 and 3 correspond to * and **, respectively.

All replicas showed similar unfolding profiles (*SI Appendix*, Fig. S4 *A–C*) where the first unfolding event always corresponded to detachment of the N-terminal helix, resulting in the first peak and subsequent force drop in simulated FECs (Fig. 3 *A,B* and *SI Appendix*, Fig. S5 *A,B*, events labeled 1* and 2). Unfolding continued from the C-terminal end of the protein (Fig. 3 *C*) until ∼34 nm of extension. At this point, the second N-terminal helix is detached and unfolded corresponding to another major peak in the FEC (Fig. 3 *A,B* and *SI Appendix*, Fig. S5 *A,B*, events labeled 3** and 4). These two peaks were separated by 24 nm in extension, which is close to the Δ*L*_*0*_ of one of the most common intermediates observed in the AFM experiments (∼27 nm) (*SI Appendix*, Fig. S6). We note that extension approximately equals *L*_0_ at the elevated rupture forces in the SMD trajectories (>200 pN). Energy barriers in unfolding pathways can be broad (64) and their crossing is a stochastic, thermally driven process. Therefore, the height and precise position of peaks in the FEC varies in AFM experiment and SMD simulation trajectories. Nevertheless, there is general agreement between the experimental Δ*L*_*0*_ and the extension difference in the simulated FECs (*SI Appendix*, Fig. S6). This suggests that the SMD simulations have identified the molecular events associated with unfolding in the AFM experiments. Importantly, these simulations show that NleC unfolding initiates at the N-terminus (Fig. 3 *B*).

All-atom SMD simulations were also carried out for protealysin (1–341 aa) at a pulling speed of 2.73 Å/ns in triplicate. All replicas showed a similar unfolding pathway (*SI Appendix*, Fig. S4 *D–F*). In contrast to NleC, the first unfolding event of protealysin was always the detachment of the C-terminal helix, which corresponds to a major peak (and drop) in the FEC (Fig. 3 *D,E* and *SI Appendix*, Fig. S5 *C,D*, events labeled 1* and 2). Protealysin then unfolded sequentially from the C-terminus (Fig. 3 *F*). Another force peak and force drop was observed at ∼24 nm extension, which corresponds to detachment and unfolding of a helix-turn-helix motif that is C-terminal of the conserved zincin fold of protealysin (Fig. 3 *D,E* and *SI Appendix*, S5 *C,D*, events labeled 3** and 4). The peaks are separated by 22 nm in extension, in good agreement with the Δ*L*_*0*_ for the most common intermediate observed in the AFM experiments (Δ*L*_*0*_ ∼21 nm) (*SI Appendix*, Fig. S7). This agreement supports the usefulness of the SMD trajectories to interpret the AFM unfolding experiments. For protealysin, unfolding proceeds from the C-terminus. This implies that the N-terminal domain of protealysin is mechanically robust compared to its C-terminus such that detaching it from the intact structure would require even higher forces.

### Biological Implications

T3SS effectors are synthesized and folded in the bacterial cytosol prior to secretion. Some effectors, such as SopE2 and SptP, have an N-terminal (∼100 aa) chaperone binding domain. Specialized chaperones bind these regions and keep the N-terminus in extended, loosely folded conformations whereas the C-terminal effector domains are well folded (65-67). Similarly, chaperone-independent effectors such as NleC are well folded in the bacterial cytosol. Indeed, expression of *Citrobacter rodentium* NleC together with its eukaryotic substrate NF-*κ*B p65 resulted in intra-bacterial cleavage of p65 demonstrating that NleC was enzymatically active and thus folded (68).

As the lumen of the T3SS needle is narrow (∼2 nm), effector proteins must be unfolded for secretion. However, the unfoldase is relatively weak and unable to unfold proteins that have been shown to be mechanically stable (*e.g*. GFP, ubiquitin and DHFR). Previous work established that effectors have typical thermodynamic stabilities, and thus thermodynamic stability does not correlate with secretion competency (29). Conversely, mechanical stability was negatively correlated with secretion competency. Specifically, *Salmonella* effectors SopE2 and SptP were shown to be mechanically labile (unfolding at low forces) and compliant (long Δ*x*^‡^), while GFP, ubiquitin and DHFR were mechanically robust (unfolding at higher forces) and brittle (short Δ*x*^‡^) (29).

Here we showed that the enterohemorrhagic *E. coli* effector NleC is also mechanically labile and unusually compliant, further cementing these features as the mechanical fingerprint that defines T3SS effectors. NleC belongs to the MA clan of zinc-dependent proteases containing a conserved zincin fold. Notably, the NleC non-effector homolog protealysin, another MA clan zincin, was more mechanically stable, requiring higher forces for unfolding, as well as brittle (short Δ*x*^‡^). These observations support a model that effector NleC diverged from other zincins to evolve structural features that would confer mechanical lability for efficient unfolding by the weak T3SS unfoldase. Such a particular evolutionary pressure is consistent with the sequence divergence observed in T3SS effectors, which in the case of NleC results in less than 15% sequence identity to any other non-effector zincin.

SMD simulations of NleC and protealysin provide molecular insight into the unfolding pathways under force. The pulling geometry in these simulations across the N- and C-termini is identical to that in AFM experiments. Under these conditions, NleC consistently unfolded from the N-terminus, indicating that this is the part of the protein easiest to unfold by force. This is relevant for the T3SS unfoldase that may work by pulling effectors from their N-termini through a narrow pore to force their unfolding end to end. For NleC, such an unfoldase would unfold the protein from its weakest point. Conversely, protealysin consistently unfolded from its C-terminus implying that its N-terminus is more resistant to unfolding. In the AFM geometry, protealysin required higher forces than NleC to be unfolded, but even higher forces would be required to unfold protealysin from its N-terminus by the T3SS unfoldase during secretion.

This observation leads to the hypothesis that NleC evolved an N-terminal structure divergent from other zincins to be mechanically labile. In Fig. 4, we compare the three-dimensional structures of NleC and five representative zincins with the conserved zincin fold composed of a three-helix and β-sheet. The segment N-terminal to the conserved fold is highlighted. Notably, NleC has two N-terminal helices that pack against the conserved zincin fold. On the other hand, all other zincins have N-terminal structures that add one or more β-strands to the conserved β-sheets. As α-helices are mechanically labile compared to β-sheets (54), this is consistent with the model that NleC evolved a divergent N-terminal structure that is mechanically labile.

**Figure 4:**
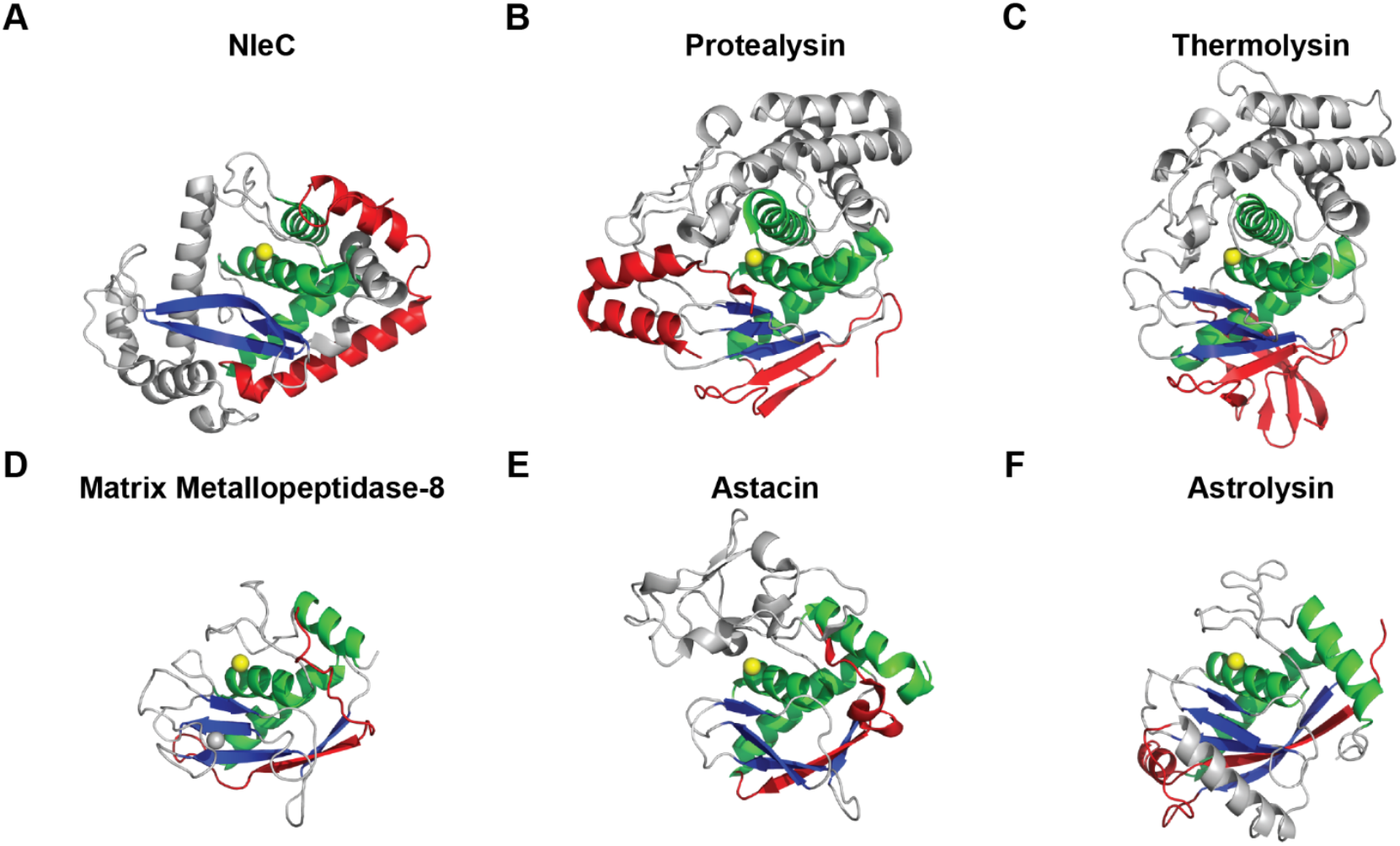
Divergent N-terminal region of representative Zincins. Zincins’ (A–H) folds (grey) are defined by a conserved three helix bundle (green) and β-sheet (blue) surrounding the catalytic zinc (yellow). The N-terminal peripheral structures (red) to the catalytic core are divergent. (A) NleC (M85 family; PDB 4Q3J); (B) protealysin (M4 family; PDB 2VQX); (C) thermolysin (M4 family; PDB 1TLP); (D) matrix metalloprotease-8 (M10A family; PDB 1KBC); (E) astacin (M12 family; PDB 1AST); (F) astrolysin (M12 family; PDB 1DTH)

In conclusion, our results indicate that effector NleC evolved from non-effector homolog protealysin to be mechanically labile and compliant to enable its unfolding for secretion. Future secretion kinetic assays could determine whether the differences in mechanical properties of NleC and protealysin render unfolding rate-limiting, thereby establishing mechanical lability as a determinant of secretion. Moreover, NleC unfolding initiates from the N-terminus, and structural comparisons to other zincins indicate that NleC’s divergent and α-helical N-terminal features confer mechanical lability. Engineering the N-terminus to produce a mechanically stabilized NleC variant would therefore provide valuable insight into how mechanical lability is encoded.

## Materials and Methods

### Functionalization of Cantilevers and Surfaces for AFM

Cantilevers (BioLever Mini, Olympus) were modified with a focused-ion beam (FIB) to improve force precision, stability and time resolution according to established methods (42, 69). Cantilevers and surfaces were then silanized and PEGylated using one of two methods described below.

In the first approach, maleimide-functionalized surfaces and cantilevers were created using established methods (70). In the second approach, surfaces were reacted with 3-aminopropyldimethylethoxysilane [APDMES, Gelest, 1.8% (v/v)] in ethanol) for 1 h at room temperature whereas cantilevers were reacted with APDMES [50% (v/v)] for 15 min at room temperature. Surfaces and cantilevers were then washed and incubated in an oven under vacuum at 80 °C for 1 h.

Amine-functionalized cantilevers and surfaces then proceeded to PEGylation where they were incubated with borate buffer (50 mM Na_2_B_4_O_7_ pH 8.5) for 1 h at room temperature. After removing excess buffer, a maleimide-PEG_12_-TFP ester [Vector Laboratories, 25 mM in 25% DMSO (v/v): 75% (v/v) borate buffer] solution was reacted with surfaces or cantilevers for 1 h at room temperature. They were then thoroughly rinsed with ultrapure water.

We immediately followed PEGylation by chemically coupling the maleimide of surfaces and cantilevers to CoA (1 mM in 50 mM sodium phosphate pH 7.2, 50 mM NaCl, 10 mM EDTA) for 1 h at room temperature or overnight at 4 °C. We coupled polyproteins and cohesin to CoA-surfaces and cantilevers, respectively, by sfp phosphopantetheinyl transferase (sfp) (37). CoA-surfaces were reacted with polyprotein (0.1–1 μM), sfp (3 μM) and MgCl_2_ (10 μM) whereas CoA-cantilevers were reacted with Cohesin (20 μM), sfp (3 μM) and MgCl_2_ (10 μM) for in AFM Buffer (25 mM HEPES, pH 7.2, 150 mM NaCl) for 1 h at room temperature or overnight at 4 °C. The surfaces and cantilevers were then thoroughly rinsed and immediately used for AFM measurements. Additional details to functionalization protocols can be found in the *SI Appendix*.

### AFM Assay and Analysis

We used a Cypher ES (Asylum Research) outfitted with a custom small spot laser (43) and temperature-controlled (25 ºC) fluid cell to collect AFM data sets. Cantilever sensitivity (V/nm) was determined by hard contact with a rigid surface and the spring constant (*k*) via the thermal method (71). To attach polyprotein to the cantilever, the cohesin-coated cantilever was gently (75–100 pN) pressed into the surface for 0.15 s. The cantilever was then retracted at a constant velocity ranging from 30–3200 nm/s and raster scanned the surface for attachments. To improve precision, AFM data sets were collected with a single cantilever (41).

The resulting FECs were digitized at 50 kHz with a low-pass filter at 25 kHz. Force was determined from the deflection of the cantilever using the calibrated cantilever spring constant and sensitivity. Extension was determined by stage movement minus cantilever deflection. We calculated the loading rate and rupture force by fitting a line to a force versus time plot immediately before an unfolding event. All FEC segments were fitted by the WLC model to determine the *L*_0_. Further details on the AFM assay and analysis of FECs can be found in the *SI Appendix*.

### Kinetic Unfolding Measurements and Analysis

Kinetic unfolding measurements were collected with an Applied PhotoPhysics ChiraScan Plus by manual mixing or outfitted with a SX-20 Stop Flow spectrophotometer. The spectrophotometer was purged with N_2_ to prevent ozone formation before use. Unfolding was initiated by the rapid mixing of buffered (SopE2: 25 mM HEPES pH 7.2, 150 mM NaCl; SptP: 10 mM Tris pH 8, 150 mM sodium Sulfate; NleC: 25 mM HEPES pH 7.2, 150 mM sodium sulfate) denaturant (urea or guanidinium hydrochloride) with SopE2, SptP or NleC to final concentrations of 0.18, 0.10 or 0.05 mg/mL, respectively. Kinetic measurements were carried out at various denaturant concentrations for 5–120 s at 222 (SptP, SopE2) or 225 nm (NleC) nm. Three independent replicates were collected and fitted with a single or double exponential fit to extract the unfolding rate constants (for double exponential fits, the fast rate constant was used).

The natural logarithm of the rate of unfolding linearly varies with [D]:

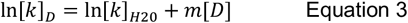

where [*k*]_D_ is the rate of unfolding in denaturant; [k]_H20_ is the rate of unfolding in water; *m* is the constant of proportionality; [*D*] is denaturant. The natural logarithm of *k* was plotted against denaturant concentrations, fitted with equation 3 and 95% confidence intervals. The fit was extrapolated to determine k_0_. More detailed procedures can be found in *SI Appendix*.

### SMD simulations

SMD simulations (72, 73) were performed to describe the unfolding mechanism of NleC and protealysin. The crystal structure of NleC (PDB: 4Q3J) and an AlphaFold2 model of protealysin (derived from PDB: 2VQX) were used as the starting structures for the simulations. Protein topologies were generated using VMD’s psfgen plugin (74). The proteins were solvated and ionized using VMD’s solvation and autoionize plugs (74). The net charge of the system was neutralized using a 150 mM of NaCl. To relax the structure and prepare for SMD simulations, the proteins were minimized for 10,000 steps and equilibrated for 5 ns with a harmonic constrain with a force constant (*k*) of 1 kcal/mol/Å^2^. SMD simulations were performed once the protein structure was properly equilibrated and checked for stability. The solvent box was enlarged in the Z direction to accommodate at least 300,000 atoms, which allows for a complete unfolding of a 180 amino-acid protein. Multiple SMD simulations were performed at pulling velocities of 2.06 Å/ns (NleC) and 2.73 Å/ns (protealysin) using the end-to-end distance of the protein as the reaction coordinate. In all the simulations the N-terminal residue of the protein was harmonically restrained to its initial position and a moving restraint with *k* = 10 kcal/mol/Å^2^ was applied to the C-terminal residue of the protein. The applied force on the harmonic spring was monitored during the entire SMD simulation and plotted against the end-to-end distance. All MD simulations were performed using NAMD3 (75, 76). The system was described by the CHARMM36m force field (77, 78) for proteins and TIP3P for water (79). Long-range electrostatic forces were calculated using particle-mesh Ewald (PME) (80), and a distance cutoff of 12 Å was used to model the short-range, non-bonded interactions. The integration time was 2 fs, and the temperature was maintained at 310 K by Langevin thermostat with a damping coefficient of 1 ps^-1^. All simulations were run as NPT ensembles.

## Acknowledgements

We thank the Biochemistry Department Shared Instruments Pool at CU Boulder (RRID: SCR_018986) for use of its general infrastructure (centrifuges, Avestin C3 homogenizer, Applied PhotoPhysics ChiraScan Plus CD and stopped flow modules). We are immensely grateful for the support provided by SIP staff members Dr. Annette Erbse and Emily Proksch. This work was supported in part by NSF grant 2145848 to MCS and ET, NSF Grant MCB-2139572 and NIST funding to TTP, NIH grant R24GM145965 to ET and NIH Molecular Biophysics T32 Training Grant GM065103 supporting KED.

## References

1. D. M. Anderson, O. Schneewind, A mRNA signal for the type III secretion of Yop proteins by Yersinia enterocolitica. Science 278, 1140–1143 (1997).

2. R. Arnold et al., Sequence-Based Prediction of Type III Secreted Proteins. PLoS Pathog 5, e1000376 (2009).

3. S. C. Birtalan, R. M. Phillips, P. Ghosh, Three-dimensional secretion signals in chaperone-effector complexes of bacterial pathogens. Mol. Cell 9, 971–980 (2002).

4. Y. Luo et al., Structural and biochemical characterization of the type III secretion chaperones CesT and SigE. Nat Struct Biol 8, 1031–1036 (2001).

5. T. Michiels, P. Wattiau, R. Brasseur, J. M. Ruysschaert, G. Cornelis, Secretion of Yop proteins by Yersiniae. Infect Immun 58, 2840–2849 (1990).

6. H. Russmann, T. Kubori, J. Sauer, J. E. Galan, Molecular and functional analysis of the type III secretion signal of the Salmonella enterica InvJ protein. Mol Microbiol 46, 769–779 (2002).

7. Y. Akeda, J. E. Galán, Chaperone release and unfolding of substrates in type III secretion. Nature 437, 911–915 (2005).

8. J. E. Galan, G. Waksman, Protein-Injection Machines in Bacteria. Cell 172, 1306–1318 (2018).

9. P. C. Lee, A. Rietsch, Fueling type III secretion. Trends Microbiol 23, 296–300 (2015).

10. A. Alonso et al., Lck dephosphorylation at Tyr-394 and inhibition of T cell antigen receptor signaling by Yersinia phosphatase YopH. J Biol Chem 279, 4922–4928 (2004).

11. D. S. Black, J. B. Bliska, Identification of p130Cas as a substrate of Yersinia YopH (Yop51), a bacterial protein tyrosine phosphatase that translocates into mammalian cells and targets focal adhesions. EMBO J 16, 2730–2744 (1997).

12. S. Mühlen, M.-H. Ruchaud-Sparagano, B. Kenny, Proteasome-independent degradation of canonical NFkappaB complex components by the NleC protein of pathogenic Escherichia coli. J. Biol. Chem. 286, 5100–5107 (2011).

13. H. Yen et al., NleC, a Type III Secretion Protease, Compromises NF-*κ*B Activation by Targeting p65/RelA. PLoS Path. 6, e1001231 (2010).

14. J. S. Pearson, P. Riedmaier, O. Marches, G. Frankel, E. L. Hartland, A type III effector protease NleC from enteropathogenic Escherichia coli targets NF-*κ*B for degradation. Mol. Microbiol. 80, 219–230 (2011).

15. D. Tezcan-Merdol, L. Engstrand, M. Rhen, Salmonella enterica SpvB-mediated ADP-ribosylation as an activator for host cell actin degradation. Int. J. Med. Microbiol. 295, 201–212 (2005).

16. Y. Fu, J. E. Galán, A salmonella protein antagonizes Rac-1 and Cdc42 to mediate host-cell recovery after bacterial invasion. Nature 401, 293–297 (1999).

17. M. C. Schlumberger, W. D. Hardt, Triggered phagocytosis by Salmonella: bacterial molecular mimicry of RhoGTPase activation/deactivation. Curr. Top. Microbiol. Immunol. 291, 29–42 (2005).

18. A. Peth, J. A. Nathan, A. L. Goldberg, The ATP costs and time required to degrade ubiquitinated proteins by the 26 S proteasome. J Biol Chem 288, 29215–29222 (2013).

19. S. M. Doyle, O. Genest, S. Wickner, Protein rescue from aggregates by powerful molecular chaperone machines. Nature Reviews Molecular Cell Biology 14, 617–629 (2013).

20. A. O. Olivares, H. C. Kotamarthi, B. J. Stein, R. T. Sauer, T. A. Baker, Effect of directional pulling on mechanical protein degradation by ATP-dependent proteolytic machines. Proc Natl Acad Sci U S A 114, E6306–E6313 (2017).

21. A. R. Nager, T. A. Baker, R. T. Sauer, Stepwise unfolding of a beta barrel protein by the AAA+ ClpXP protease. J Mol Biol 413, 4–16 (2011).

22. R. Zarivach, M. Vuckovic, W. Y. Deng, B. B. Finlay, N. C. J. Strynadka, Structural analysis of a prototypical ATPase from the type III secretion system. Nat. Struct. Mol. Biol. 14, 131–137 (2007).

23. D. D. Majewski et al., Cryo-EM structure of the homohexameric T3SS ATPase-central stalk complex reveals rotary ATPase-like asymmetry. Nat Commun 10, 626 (2019).

24. V. T. Lee, O. Schneewind, Yop fusions to tightly folded protein domains and their effects on Yersinia enterocolitica type III secretion. J Bacteriol 184, 3740–3745 (2002).

25. J. A. Sorg, N. C. Miller, M. M. Marketon, O. Schneewind, Rejection of impassable substrates by Yersinia type III secretion machines. J Bacteriol 187, 7090–7102 (2005).

26. J. Radics, L. Konigsmaier, T. C. Marlovits, Structure of a pathogenic type 3 secretion system in action. Nat Struct Mol Biol 21, 82–87 (2014).

27. K. Dohlich, A. B. Zumsteg, C. Goosmann, M. Kolbe, A substrate-fusion protein is trapped inside the Type III Secretion System channel in Shigella flexneri. PLoS Pathog 10, e1003881 (2014).

28. J. E. Dawson, L. K. Nicholson, Folding kinetics and thermodynamics of Pseudomonas syringae effector protein AvrPto provide insight into translocation via the type III secretion system. Protein Sci 17, 1109–1119 (2008).

29. M. A. LeBlanc, M. R. Fink, T. T. Perkins, M. C. Sousa, Type III secretion system effector proteins are mechanically labile. Proc Natl Acad Sci U S A 118 (2021).

30. K. Baruch et al., Metalloprotease type III effectors that specifically cleave JNK and NF-kB. EMBO J, 1–11 (2010).

31. M. M. Turco, M. C. Sousa, The structure and specificity of the type III secretion system effector NleC suggest a DNA mimicry mechanism of substrate recognition. Biochemistry 53, 5131–5139 (2014).

32. R. Arnold et al., Sequence-based prediction of type III secreted proteins. PLoS Pathog 5, e1000376 (2009).

33. N. D. Rawlings et al., The MEROPS database of proteolytic enzymes, their substrates and inhibitors in 2017 and a comparison with peptidases in the PANTHER database. Nucleic Acids Res 46, D624–D632 (2018).

34. A. G. Murzin, S. E. Brenner, T. Hubbard, C. Chothia, SCOP: A structural classification of proteins database for the investigation of sequences and structures. J Mol Biol 247, 536–540 (1995).

35. M. Punta et al., The Pfam protein families database. Nucleic Acids Res 40, D290–D301 (2011).

36. I. V. Demidyuk et al., Crystal structure of the protealysin precursor: insights into propeptide function. J Biol Chem 285, 2003–2013 (2010).

37. J. Yin, A. J. Lin, D. E. Golan, C. T. Walsh, Site-specific protein labeling by Sfp phosphopantetheinyl transferase. Nat Protoc 1, 280–285 (2006).

38. M. Schlierf, F. Berkemeier, M. Rief, Direct observation of active protein folding using lock-in force spectroscopy. Biophys J 93, 3989–3998 (2007).

39. C. Schoeler et al., Mapping Mechanical Force Propagation through Biomolecular Complexes. Nano Lett. 15, 7370–7376 (2015).

40. S. W. Stahl et al., Single-molecule dissection of the high-affinity cohesin-dockerin complex. Proc Natl Acad Sci U S A 109, 20431–20436 (2012).

41. M. Otten et al., From genes to protein mechanics on a chip. Nat Methods 11, 1127–1130 (2014).

42. M. S. Bull, R. M. Sullan, H. Li, T. T. Perkins, Improved single molecule force spectroscopy using micromachined cantilevers. ACS Nano 8, 4984–4995 (2014).

43. D. T. Edwards et al., Optimizing 1-mu s-Resolution Single-Molecule Force Spectroscopy on a Commercial Atomic Force Microscope. Nano Lett. 15, 7091–7098 (2015).

44. D. T. Edwards, T. T. Perkins, Optimizing force spectroscopy by modifying commercial cantilevers: Improved stability, precision, and temporal resolution. Journal of Structural Biology 197, 13–25 (2017).

45. C. Bustamante, J. F. Marko, E. D. Siggia, S. Smith, Entropic elasticity of lambda-phage DNA. Science 265, 1599–1600 (1994).

46. M. Rief, M. Gautel, F. Oesterhelt, J. M. Fernandez, H. E. Gaub, Reversible unfolding of individual titin immunoglobulin domains by AFM. Science 276, 1109–1112 (1997).

47. H. Dietz, M. Rief, Exploring the energy landscape of GFP by single-molecule mechanical experiments. P Natl Acad Sci USA 101, 16192–16197 (2004).

48. B. Yang, Z. Liu, H. Liu, M. A. Nash, Next Generation Methods for Single-Molecule Force Spectroscopy on Polyproteins and Receptor-Ligand Complexes. Front Mol Biosci 7, 85 (2020).

49. Y. Cao, H. Li, Polyprotein of GB1 is an ideal artificial elastomeric protein. Nat Mater 6, 109–114 (2007).

50. E. Evans, K. Ritchie, Strength of a weak bond connecting flexible polymer chains. Biophys J 76, 2439–2447 (1999).

51. E. Evans, K. Ritchie, Dynamic strength of molecular adhesion bonds. Biophys. J. 72, 1541–1555 (1997).

52. C. L. Chyan et al., Reversible mechanical unfolding of single ubiquitin molecules. Biophys J 87, 3995–4006 (2004).

53. S. R. Ainavarapu, L. Li, C. L. Badilla, J. M. Fernandez, Ligand binding modulates the mechanical stability of dihydrofolate reductase. Biophys J 89, 3337–3344 (2005).

54. T. Hoffmann, K. M. Tych, M. L. Hughes, D. J. Brockwell, L. Dougan, Towards design principles for determining the mechanical stability of proteins. Phys Chem Chem Phys 15, 15767–15780 (2013).

55. G. I. Bell, Models for the specific adhesion of cells to cells. Science 200, 618–627 (1978).

56. M. T. Woodside, S. M. Block, Reconstructing folding energy landscapes by single-molecule force spectroscopy. Annu Rev Biophys 43, 19–39 (2014).

57. S. O. Garbuzynskiy, D. N. Ivankov, N. S. Bogatyreva, A. V. Finkelstein, Golden triangle for folding rates of globular proteins. Proc Natl Acad Sci U S A 110, 147–150 (2013).

58. A. V. Glyakina, O. V. Galzitskaya, How Quickly Do Proteins Fold and Unfold, and What Structural Parameters Correlate with These Values? Biomolecules 10 (2020).

59. M. Sotomayor, K. Schulten, Single-molecule experiments in vitro and in silico. Science 316, 1144–1148 (2007).

60. D. Sharma et al., Single-molecule force spectroscopy reveals a mechanically stable protein fold and the rational tuning of its mechanical stability. Proc Natl Acad Sci U S A 104, 9278–9283 (2007).

61. J. Schonfelder, R. Perez-Jimenez, V. Munoz, A simple two-state protein unfolds mechanically via multiple heterogeneous pathways at single-molecule resolution. Nat Commun 7, 11777 (2016).

62. T. Verdorfer et al., Combining in Vitro and in Silico Single Molecule Force Spectroscopy to Characterize and Tune Cellulosomal Scaffoldin Mechanics. J Am Chem Soc 10.1021/jacs.7b07574 (2017).

63. B. Yang et al., Engineering the Mechanical Stability of a Therapeutic Complex between Affibody and Programmed Death-Ligand 1 by Anchor Point Selection. ACS Nano 18, 31912–31922 (2024).

64. S. P. Ng et al., Mechanical unfolding of TNfn3: the unfolding pathway of a fnIII domain probed by protein engineering, AFM and MD simulation. J Mol Biol 350, 776–789 (2005).

65. M. Iimori et al., Structural basis of effector recognition by the T3SS chaperone VecA from Vibrio parahaemolyticus. Biochem Biophys Res Commun 776, 152190 (2025).

66. M. Lilic, M. Vujanac, C. E. Stebbins, A Common Structural Motif in the Binding of Virulence Factors to Bacterial Secretion Chaperones. Mol. Cell 21, 653–664 (2006).

67. F. D. Schubot et al., Three-dimensional structure of a macromolecular assembly that regulates type III secretion in Yersinia pestis. J Mol Biol 346, 1147–1161 (2005).

68. M. K. Hasan, S. El Qaidi, P. R. Hardwidge, The T3SS Effector Protease NleC Is Active within Citrobacter rodentium. Pathogens 10 (2021).

69. J. K. Faulk, D. T. Edwards, M. S. Bull, T. T. Perkins, Improved Force Spectroscopy Using Focused-Ion-Beam-Modified Cantilevers. Methods Enzymol 582, 321–351 (2017).

70. R. Walder et al., Rapid Characterization of a Mechanically Labile alpha-Helical Protein Enabled by Efficient Site-Specific Bioconjugation. J Am Chem Soc 139, 9867–9875 (2017).

71. J. E. Sader, J. W. M. Chon, P. Mulvaney, Calibration of rectangular atomic force microscope cantilevers. Rev Sci Instrum 70, 3967–3969 (1999).

72. B. Isralewitz, M. Gao, K. Schulten, Steered molecular dynamics and mechanical functions of proteins. Curr Opin Struct Biol 11, 224–230 (2001).

73. S. Izrailev, S. Stepaniants, M. Balsera, Y. Oono, K. Schulten, Molecular dynamics study of unbinding of the avidin-biotin complex. Biophys J 72, 1568–1581 (1997).

74. W. Humphrey, A. Dalke, K. Schulten, VMD: visual molecular dynamics. J Mol Graph 14, 33-38, 27-38 (1996).

75. J. C. Phillips et al., Scalable molecular dynamics with NAMD. J Comput Chem 26, 1781–1802 (2005).

76. J. C. Phillips et al., Scalable molecular dynamics on CPU and GPU architectures with NAMD. J Chem Phys 153, 044130 (2020).

77. R. B. Best et al., Optimization of the additive CHARMM all-atom protein force field targeting improved sampling of the backbone phi, psi and side-chain chi(1) and chi(2) dihedral angles. J Chem Theory Comput 8, 3257–3273 (2012).

78. J. Huang et al., CHARMM36m: an improved force field for folded and intrinsically disordered proteins. Nat Methods 14, 71–73 (2017).

79. W. L. Jorgensen, J. Chandrasekhar, J. D. Madura, R. W. Impey, M. L. Klein, Comparison of Simple Potential Functions for Simulating Liquid Water. J Chem Phys 79, 926–935 (1983).

80. U. Essmann et al., A Smooth Particle Mesh Ewald Method. J Chem Phys 103, 8577–8593 (1995).

