## Supplementary Appendix for "A Type III secretion system effector evolved to be mechanically labile and initiate unfolding from the N-terminus"

### Supplementary Materials

#### Materials and Methods

##### AFM constructs and cloning

The gene encoding the catalytic domain NleC (residues 22–280) was amplified by PCR using primers U2418 and L2418 (*SI Appendix*, Table S2). The gene encoding the full-length inactive protealysin (residues 6–341, E163A) was amplified by PCR using primers U2515 and L2483. NleC gene fragments were each incorporated into plasmid pMS-1480 linearized by PCR amplification with primers U2414 and L2414 by Gibson assembly using NEBuilder HiFi DNA Assembly (New England Biolabs). The protealysin gene fragments were integrated into plasmid pMS-1711 linearized by restriction digest with ZraI also by Gibson assembly. The construct with NleC is pMS-1620 and protealysin is pMS-1696. The vectors pMS-1480 and pMS-1711 are both pET-28a based expression vectors. pMS-1480 includes a 6x His-tag for Ni-NTA affinity chromatography, a ybbR-tag that allows for covalent attachment to CoA-derivatized surfaces, well characterized 4th domain of the F-actin cross-linking filamin rod of Dictostelium discoideum (ddFLN4) as a marker protein (1) for single-molecule attachment, and Xmod-Dockerin for strong, noncovalent attachment of the polyprotein to cohesin functionalized cantilevers. pMS-1711 has all the same features as pMS-1480 except it includes a different marker protein, streptococcal B1 immunoglobulin-binding domain of protein G or GB1 (2), rather than ddFLN4.

##### Protein expression and purification for AFM

The plasmids encoding the polyproteins were transformed into E. coli BL21 (D3E) cells (GoldBio), and single colonies were grown overnight at 37 °C with shaking in LB supplemented with kanamycin. One liter LB + kanamycin cultures were inoculated [1:100 (v/v)] and grown at 37 °C with shaking. Once an $OD_{600}$ of 0.6 was reached, cultures were put on ice for 30 min and then induced with 0.4 mM IPTG at 16 °C overnight.

Cultures were pelleted at 4000 RPM with a Beckman JLA 8.100 rotor for 30 min at 4 °C, and cell pellets were resuspended with lysis buffer [50 mM Tris pH 8, 0.5 mg/mL lysozyme, EDTA-free protease inhibitor tablets (Pierce)]. Cells were lysed by sonication on ice or a C3 homogenizer. Cell debris was removed by centrifugation in a Sorvall SS-34 rotor at 14,000 x g for 30 min at 4 °C. The supernatant was filtered through 0.22 µM PVDF membrane before loaded onto a 5-mL Ni-NTA (Cytiva) column equilibrated with Buffer A (50 mM Tris pH 8, 150 mM NaCl). The column was then washed with 5 column volumes (CVs) of Buffer A + 20 mM Imidazole and eluted off the column with a linear gradient of 5 CVs of Buffer A + 25–300 mM Imidazole. Fractions containing the protein of interest were then loaded onto a size exclusion column (HiLoad 26/60 Superdex 200, Amersham Pharmacia Biotech) equilibrated with AFM buffer (25 mM HEPES pH 7.2, 150 mM NaCl, 0.5 mM TCEP). Fractions containing the protein of interest were combined, concentrated, and stored at -80 °C for later use.

##### Functionalization of cantilevers and surfaces for AFM

Cantilevers (BioLever Mini, Olympus) were modified with a focused-ion beam (FIB) according to previously described methods (3, 4). Briefly, cantilevers were modified to remove large sections to reduce hydrodynamic drag, thinned to reduce stiffness, and chemically etched to remove the majority of the gold and chromium coatings to improve stability. We initially cleaned glass surfaces (German Glass, Electron Microscopy Sciences) for functionalization by rinsing in subsequent baths of ultrapure water, acetone, ethanol, 3 M KOH, and two additional rinses with ultrapure water. Surfaces were then dried with $N_2$ gas and irradiated with ozone for 30 min (PSD-UV8, Novascan Technologies). Cantilevers and surfaces were then silanized and PEGylated using one of two methods described below.

In the first approach, maleimide-functionalized surfaces and cantilevers were created using established methods (5). Briefly, surfaces and cantilevers were reacted with a silane-PEG-maleimide (SPM, PEG MW = 1000 Da, Nanocs, 0.15 mg/mL in toluene) at 65 °C for 3 h. After the incubation, we washed the cantilevers and surfaces sequentially in toluene, isopropanol, and two ultrapure water baths.

The second approach involved creating amine-functionalized surfaces and cantilevers prior to PEGylation. Surfaces were incubated with 3-aminopropyldimethylethoxysilane [APDMES, Gelest, 1.8% (v/v)] in ethanol) for 1 h at room temperature whereas cantilevers were incubated with APDMES [Gelest, 50% (v/v)] for 15 min at room temperature. Surfaces and cantilevers were then washed sequentially in isopropanol and ultrapure water, dried, and incubated in an oven under vacuum at 80 °C for 1 h. Amine-functionalized cantilevers and surfaces then proceeded to PEGylation where they were incubated with borate buffer (50 mM $Na_2B_4O_7$, pH 8.5) for 1 h at room temperature. A solution of maleimide-PEG$_{12}$-TFP ester [Vector Laboratories, 25 mM in 25% DMSO (v/v): 75% (v/v) borate buffer] was made and centrifuged for 10 min at 15,000 x g. After removing excess buffer, the supernatant of the maleimide-PEG$_{12}$-TFP ester solution reacted with surfaces or cantilevers reacted for 1 h at room temperature. They were then rinsed with ultrapure water three times.

We immediately followed PEGylation by chemically coupling CoA to the maleimide-functionalized surface to minimize maleimide group hydrolysis. To do so, surfaces and cantilevers were reacted with a CoA solution (1 mM in 50 mM sodium phosphate pH 7.2, 50 mM NaCl, 10 mM EDTA) for 1 h at room temperature or overnight at 4 °C. We coupled the polyproteins or cohesin to CoA surfaces or cantilevers respectively by sfp phosphopantetheinyl transferase (sfp), which covalently attaches CoA to the ybbR-tag encoded in our constructs (6). AFM polyproteins were thawed and diluted to 0.1–1 μM in AFM buffer (25 mM HEPES, pH 7.2, and 150 mM NaCl) and combined with final concentrations of 3 μM sfp and 10 μM $MgCl_2$ and incubated for 1 h at room temperature or overnight at 4 °C. Cohesin was also deposited onto CoA-functionalized cantilevers using the same procedure with final concentrations of 20 μM cohesin, 3 μM sfp and 10 μM $MgCl_2$. The surfaces and cantilevers were then thoroughly rinsed with AFM buffer and then immediately used for AFM measurements.

**AFM assay and analysis**

If the AFM pulling assay stopped yielding attachments, the cantilevers were cleaned and the attached cohesin unfolded in a urea bath (8 M in AFM buffer) for 30 min. Cohesin on the cantilever was then refolded by soaking in AFM buffer for 30 min. Sensitivity and cantilever spring constant were recalibrated prior to use. Cantilever spring constants (*k*) ranged from ~6–15 pN/nm. This cleaning procedure often rescued attachment of the cantilever to the polyprotein. A 150-nm spaced grid on the surface was sampled. AFM experiments were stopped after ~3,000–10,000 force extension curves (FECs) were collected with a single cantilever.

In a small percentage of traces, the zero-force baseline was corrected to account for experimental drift. The FEC traces were then smoothed by box-car averaging to reduce noise for visual presentation. All FEC segments were fit by the WLC model to determine the contour length ($L_0$) of peaks. Only force extension curves displaying change in contour lengths ($\Delta L_0$) within 20% of expected values (GB1 = 18 nm; ddFLN4 = 32.2 nm; NleC = 90.4 nm; protealysin = 120.9 nm) were included in analyses. The unfolding force for NleC and protealysin was determined from the first unfolding event in the FEC when unfolding intermediates were observed. Rare high force unfolding forces exceeding 2 standard deviations from the mean were excluded from analyses, as they were likely due to cantilever-surface adhesion or protein misfolding.

**Kinetic unfolding constructs and cloning**

The catalytic domain of NleC (residues 22–280) was amplified using primers U2442 and L2438 (*SI Appendix*, Table S2). pMS-984 was digested with XhoI and SfoI. The PCR product was digested with XfoI and ligated into digested pMS-984. The plasmids encoding the catalytic domains of SopE2 (residues 69–240), SptP (residues 161–543) were obtained from reference (7). The constructs encoding SopE2, SptP and NleC are as follows: pMS-1599, pMS-1597, and pMS-1623, respectively. pMS-984, a pET-28-a based expression vector, encodes a N-terminal 6x His-tag for Ni-NTA affinity purification, a N-terminal cleavable SUMO tag to increase protein solubility, and a multiple cloning site as an alternative method to clone genes of interest.

**Protein expression and purification for kinetic unfolding measurements**

The kinetic unfolding constructs were expressed and purified using the same procedure. The constructs were transformed into E. coli BL-21 (DE3) cells (GoldBio) for overexpression. Overnight cultures inoculated 1-liter cultures [1:100 (v/v)] of LB medium supplemented with Kanamycin. Cultures grew at 37 °C with shaking until $OD_{600}$ of 0.6. Cells were then put on ice for 30 min and then induced with 0.4 mM IPTG. Cells were grown overnight at 16 °C and then pelleted for 30 min at 4 °C in a Beckman JLA 8.100 rotor at 4000 RPM.

Cell pellets were resuspended with lysis buffer [50 mM Tris pH 8, 0.5 mg/ml lysozyme, and EDTA-free protease inhibitor tablets (Pierce)]. The cells were lysed with a sonicator on ice. The cell lysate was separated from cell debris by centrifugation with a Sorvall SS-34 rotor at 14,000 x g for 30 min at 4 °C. The soluble protein fraction was loaded into a 5 mL Ni-NTA column (Cytiva) equilibrated with buffer A (50 mM

Tris pH 8, 150 mM NaCl). The column was washed with Buffer A + 20 mM imidazole for 5 CV. The column was eluted with a linear gradient of Buffer A + 20-300 mM imidazole. The eluted fractions with protein were combined and 1:1000 (w/w) of His-tagged Ulp1 enzyme was added to cleave off the N-terminal SUMO and His tag. Eluted protein and His-tagged Ulp1 were dialyzed against the reaction buffer (25 mM Tris pH 8, 150 mM NaCl, 5 mM BME) with a 6-8 kDa membrane cutoff. The cleaved protein was then loaded onto a 5-mL Ni-NTA column using the same protocol except the flow-through fractions were collected as the eluted fractions contain His-tagged Ulp1, His-tagged SUMO-tag, and uncleaved protein. The cleaved protein was then loaded onto a Superdex75 column equilibrated with buffer A. The fractions containing only protein were combined and concentrated. NleC, SptP and SopE2 were stored at -80 °C.

**Kinetic unfolding measurements and analysis**

Protein stocks were thawed from -80 °C storage and filtered with a 0.1 µM PVDF membrane. A 10 M urea buffered (SopE2: 25 mM HEPES pH 7.2, 150 mM NaCl; SptP: 10 mM Tris pH 8, 150 mM sodium sulfate) solution was made and deionized with BioRad AG 501-X8 resin (50 g/L) for 1 h. The resin was removed by vacuum filtration. A 6 M guanidinium hydrochloride buffered (25 mM HEPES pH 7.2, 150 mM sodium sulfate) solution was made. The concentrations of the urea and guanidinium hydrochloride stocks were confirmed with an Abbe refractometer.

Stopped flow data for SptP and SopE2 was collected with an Applied PhotoPhysics ChiraScan Plus outfitted with a SX-20 Stop Flow spectrophotometer with a 20 µL flow cell. Before use, the spectrophotometer was thoroughly purged with nitrogen to prevent ozone formation and protect the optics. Multiple shots (5–10) of water and buffer completely purged the lines between measurements. The absorbance spectra confirmed the system was equilibrated. Unfolding was initiated by the rapid mixing of 1:1 shot of protein and buffered urea at various concentrations chosen based on the concentration range of the transition region (7) with a dead time of 1.2 ms. The final concentration of protein after mixing was 0.18 mg/mL for SopE2 and 0.1 mg/mL for SptP. Kinetic data was collected for 5–120 s, collecting 1000 times points, at 222 nm. The collection time varied so that there would be good coverage of data points in the initial region of the unfolding curve. Three independent replicates were collected for each urea concentration. The replicates were fitted with a single or double exponential fit to the extract the unfolding rate constants (for double exponential fits, the fast rate constant was used) for each urea concentration.

The natural logarithm of the rate of unfolding linearly varies with [D]:

$$\ln[k]_D = \ln[k]_{H20} + m[D] \qquad \text{Equation 3}$$

where $[k]_D$ is the rate of unfolding in denaturant; $[k]_{H20}$ is the rate of unfolding in water or intrinsic unfolding rate, $m$ is the constant of proportionality; and $[D]$ is denaturant. The natural logarithm of $k$ was plotted against urea concentrations and fitted with equation 3 and 95% confidence intervals. The fit was extrapolated to the y-intercept to determine $k_0$.

Observing NleC unfolding kinetics required guanidinium hydrochloride concentrations that could not be achieved in stopped flow due to 1:1 dilution. Thus, we opted for manual mixing which allowed higher guanidinium hydrochloride concentrations that still captured initial unfolding rates despite increased experimental dead time. For manual mixing unfolding experiments with NleC, a quartz cuvette (Hellma) with a 10-mM path length was used with an Applied PhotoPhysics ChiraScan Plus spectrophotometer. Before use, the spectrophotometer was thoroughly purged with nitrogen to prevent ozone formation and protect the optics. NleC was mixed in the cuvette with the buffered guanidinium hydrochloride to a final concentration of 0.05 mg/mL. The time from mixing to the beginning of the data collection was noted to correct for time (~15 s). Kinetic data was collected for 120–300 s, collecting 1000 times points logarithmically at 225 nm. The cuvette was cleaned with three 10M urea washes, three buffer washes, 1–2 1% Hellmanex solution washes, five ultrapure water washes, and then one 100% ethanol wash and house air to dry the cuvette in between measurements. The resulting denaturation curves were analyzed using previously described procedures.

**Full Protein Sequences.** Individual proteins are color coded in polyprotein sequences. All construct sequences encode a 6xHis tag used for affinity purification. For construct sequences used in kinetic unfolding studies (pMS-1597, pMS-1599 and pMS-1623), a vertical bar (|) indicates the cleavage site for Ulp1 that removes the SUMO-tag.

pMS 1597 – 6xHIS-SUMO-SptP

MGSSHHHHHHSSGLVPRGSASMSDSEVNQEAKPEVKPEVKPETHINLKVSDGSSEIFFKIKKTTPLRRLMEAFAKRQ
GKEMDSLRFLYDGIRIQADQTPEDLDMEDNDIIEAHREQIG|GNDVGAESKQPLLDIALKGLKRTLPQLEQMDGNSL
RENFQEMASGNGPLRSLMTNLQNLNKIPEAKQLNDYVTTLTNIQVGVARFSQWGTCGGEVERWVDKASTHELTQAVK
KIHVIAKELKNVTAELEKIEAGAPMPQTMSGPTLGLARFAVSSIPINQQTQVKLSDGMPVPVNTLTFDGKPVALAGS
YPKNTPDALEAHMKMLLEKECSCLVVLTSEDQMQAKQLPPYFRGSYTFGEVHTNSQKVSSASQGEAIDQYNMQLSCG
EKRYTIPVLHVKNWPDHQPLPSTDQLEYLADRVKNSNQNGAPGRSSSDKHLPMIHCLGGVGRTGTMAAALVLKDNPH
SNLEQVRADFRDSRNNRMLEDASQFVQLKAMQAQLLMTTAS

pMS 1599 –6xHIS-SUMO-SopE2

MGSSHHHHHHSSGLVPRGSASMSDSEVNQEAKPEVKPEVKPETHINLKVSDGSSEIFFKIKKTTPLRRLMEAFAKRQ
GKEMDSLRFLYDGIRIQADQTPEDLDMEDNDIIEAHREQIG|GEGRAVLTSKTVKDFMLQKLNSLDIKGNASKDPAY
ARQTCEAILSAVYSNNKDQCCKLLISKGVSITPFLKEIGEAAQNAGLPGEIKNGVFTPGGAGANPFVVPLIASASIK
YPHMFINHNQQVSFKAYAEKIVMKEVTPLFNKGTMPTPQQFQLTIENIANKYLQNAS

pMS-1620 ybbR-6xHIS-ddFLN4-NleC-Xmod-Dockerin

MDSLEFIASKLAHHHHHHGSADPEKSYAEGPGLDGGESFQPSKFKIHAVDPDGVHRTDGGDGFVVTIEGPAPVDPVM
VDNGDGTYDVEFEPKEAGDYVINLTLDGDNVNGFPKTVTVKPAPGSGSGIAPNRAENAYADYVLDIGKRIPLSAADL
SNVYESVIRAVHDSRSRLIDQHTVDMIGNTVLDALSRSQTFRDAVSYGIHNEKVHIGCIKYRNEYELNEESSVKIDD
IQSLTCNELYEYDVGQEPIFPICEAGENDNEEPYVSFSVAPDTDSYEMPSWQEGLIHEIIHHVTGSSDPSGDSNIEL
GPTEILARRVAQELGWSVPDFKGYAEPEREAHLRLRNLNALRQAAMRHEENERAFFERLGTISDRYEASPDFTEYSG
SVVPNTVTSAVKTQYVEIESVDGFYFNTEDKFDTAQIKKAVLHTVYNEGYTGDDGVAVVLREYESEPVDITAELTFG
DATPANTYKAVENKFDYEIPVYYNNATLKDAEGNDATVTVYIGLKGDTDLNNIVDGRDATATLTYYAATSTDGKDAT
TVALSPSTLVGGNPESVYDDFSAFLSDVKVDAGKELTRFAKKAERLIDGRDASSILTFYTKSSVDQYKDMAANEPNK
LWDIVTGDAEEE

pMS-1623–6xHIS-SUMO-NleC

MGSSHHHHHHSSGLVPRGSASMSDSEVNQEAKPEVKPEVKPETHINLKVSDGSSEIFFKIKKTTPLRRLMEAFAKRQ
GKEMDSLRFLYDGIRIQADQTPEDLDMEDNDIIEAHREQIG|GIAPNRAENAYADYVLDIGKRIPLSAADLSNVYES
VIRAVHDSRSRLIDQHTVDMIGNTVLDALSRSQTFRDAVSYGIHNEKVHIGCIKYRNEYELNEESSVKIDDIQSLTC
NELYEYDVGQEPIFPICEAGENDNEEPYVSFSVAPDTDSYEMPSWQEGLIHEIIHHVTGSSDPSGDSNIELGPTEIL
ARRVAQELGWSVPDFKGYAEPEREAHLRLRNLNALRQAAMRHEENERAFFERLGTISDRYEASPDFTEYS

pMS-1696 ybbR-6xHIS-GB1-GB1-Protealysin-Xmod-Dockerin

MDSLEFIASKLAHHHHHHGSDTYKLILNGKTLKGETTTEAVDAATAEKVFKQYANDNGVDGEWTYDDATKTFTVTER
SDTYKLILNGKTLKGETTTEAVDAATAEKVFKQYANDNGVDGEWTYDDATKTFTVTEGSGSGPTLTARSVIPPYMLR
RIIEHGSLPQRDCALHTLNHVQSLLGNKPLRAPGAKTSTGGEVIRDIFDAENGTQLPGKQVRNEGQASNHDVAVDEA
YDYLGVTYDFFWQAFKRNSLDNQGLPLTGSVHYGKEYQNAFWNGQQMVFGDGDGEIFNRFTIAIDVVGHALAHGVTE
SEAGLIYFQQAGALNESLSDVFGSLVKQFHLKQTADKADWLIGEGLLAKGINGKGLRSMSAPGTAYNDPLLGKDPQP
ADMKDYIQTKEDNGGVHLNSGIPNRAFYLAATALGGFAWEKAGYIWYDTLCDKTLPQDADFATFARTTVKHAKQRFD
SKVADKVQQAWHQVGVASGSVVPNTVTSAVKTQYVEIESVDGFYFNTEDKFDTAQIKKAVLHTVYNEGYTGDDGVAV
VLREYESEPVDITAELTFGDATPANTYKAVENKFDYEIPVYYNNATLKDAEGNDATVTVYIGLKGDTDLNNIVDGRD
ATATLTYYAATSTDGKDATTVALSPSTLVGGNPESVYDDFSAFLSDVKVDAGKELTRFAKKAERLIDGRDASSILTF
YTKSSVDQYKDMAANEPNKLWDIVTGDAEEE

pHG 0002– CohesinIII – 6xHIS-ybbR

MALTDRGMTYDLDPKDGSSAATKPVLEVTKKVFDTAADAAGQTVTVEFKVSGAEGKYATTGYHIYWDERLEVVATKT
GAYAKKGAALEDSSLAKAENNGNGVFVASGADDDFGADGVMWTVELKVPADAKAGDVYPIDVAYQWDPSKGDLFTDN
KDSAQGKLMQAYFFTQGIKSSSNPSTDEYLVKANATYADGYIAIKAGEPHHHHHHDSLEFIASKLA

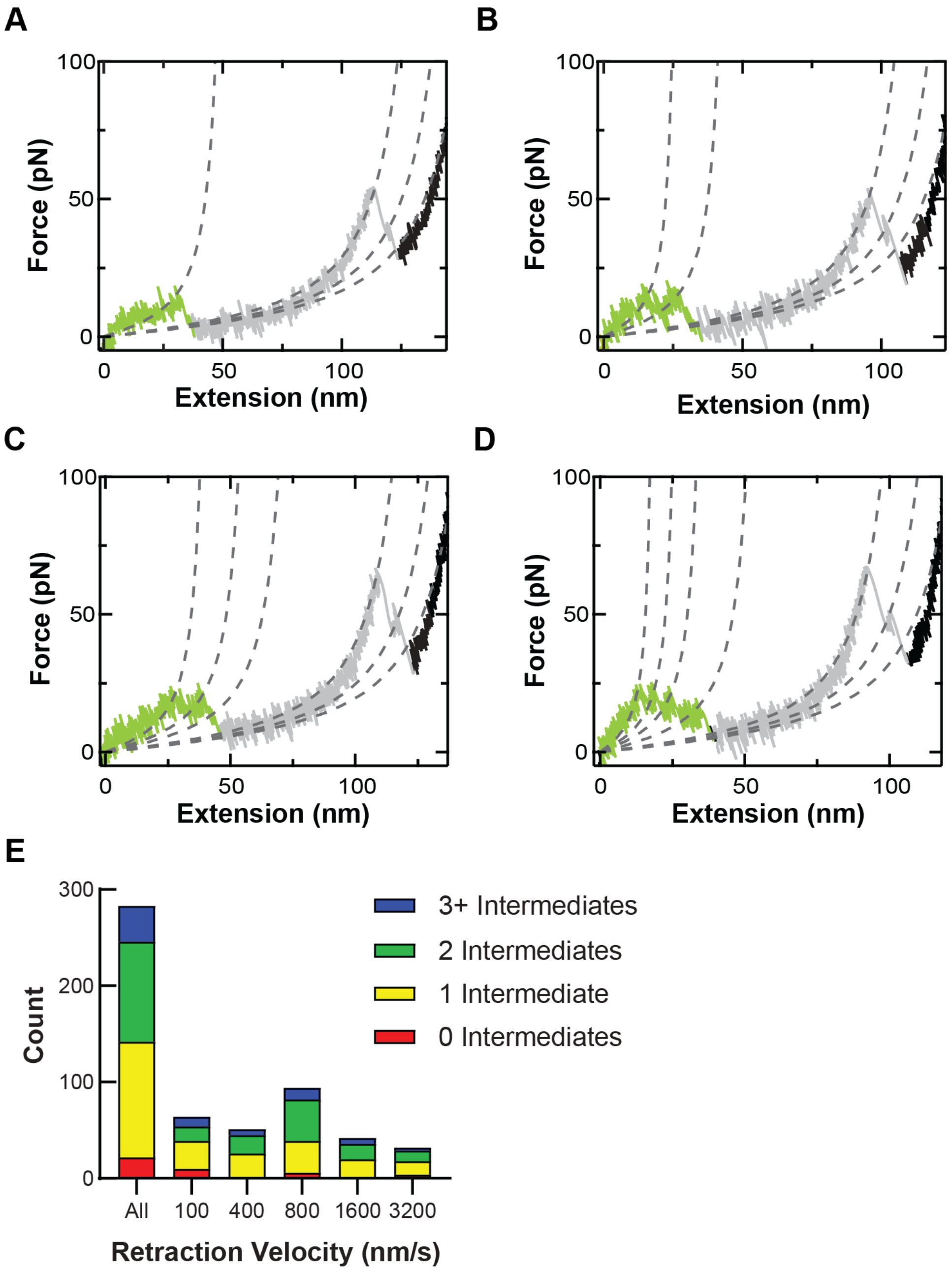


**Figure S1.** NleC preferentially unfolds through intermediates in AFM assays (A–D) FEC curves were fitted with WLC models (dashed lines) collected at $v$ = 800 nm/s depicting NleC (green) unfolding at through none (A), one (B), two (C), or three (D) intermediates. Following NleC unfolding, ddFLN4 (grey), the marker domain, unfolded. (E) Distribution of NleC unfolding through none (red), one (yellow), two (green), or multiple (blue) intermediates by retraction velocity. Bar graphs generated from N = 283, 64, 51, 94, 42, and 32 (left to right) unfolding events for the set of pulling velocities studied (100, 400, 800, 1600, and 3200 nm/s respectively).

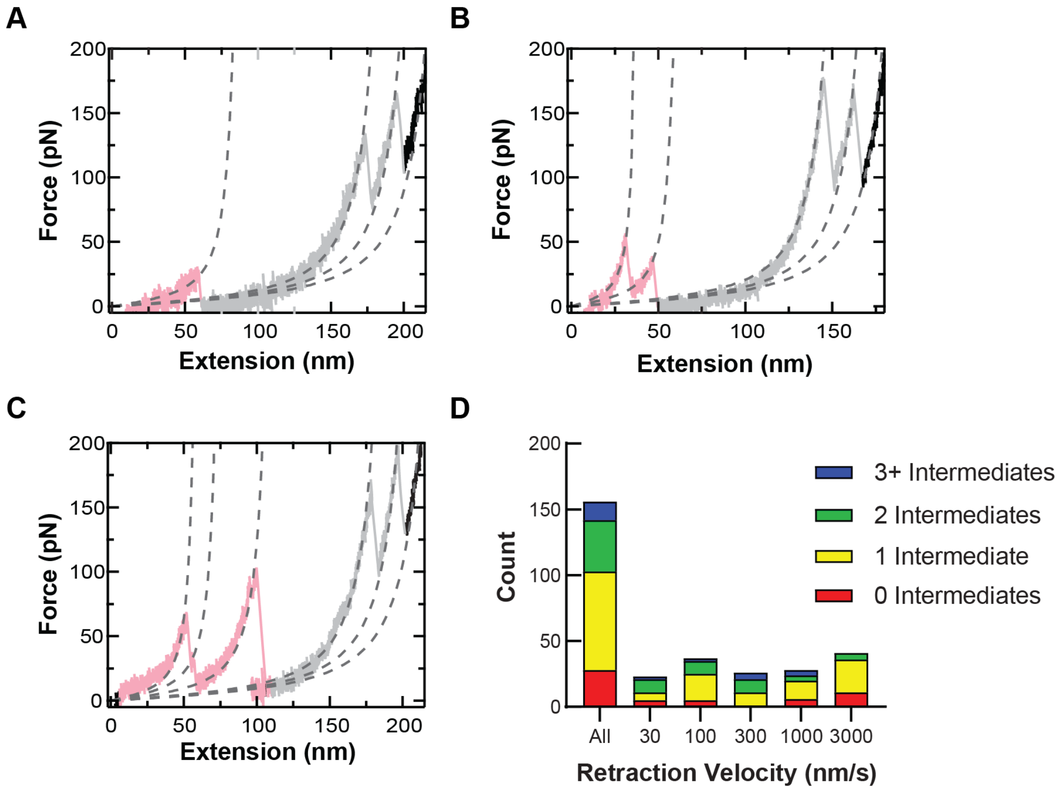


**Figure S2.** Protealysin preferentially unfolds through intermediates in AFM assays (A–C) FEC curves were fitted with WLC models (dashed lines) depicting protealysin (pink) unfolding through none (A; *v* = 1000 nm/s), one (B; *v* = 100 nm/s), or two (C; *v* = 100 nm/s). Following protealysin unfolding, two copies of the marker domain GB1 (grey) unfolded. (D) Distribution of protealysin unfolding through none (red), one (yellow), two (green), or rare multiple (blue) intermediates by retraction velocity. Bar graphs generated from N = 156, 23, 37, 26, 28, and 42 (left to right) unfolding events for *v* = all pulling velocities studied, 30, 100, 300, 1000, and 3000 nm/s respectively.

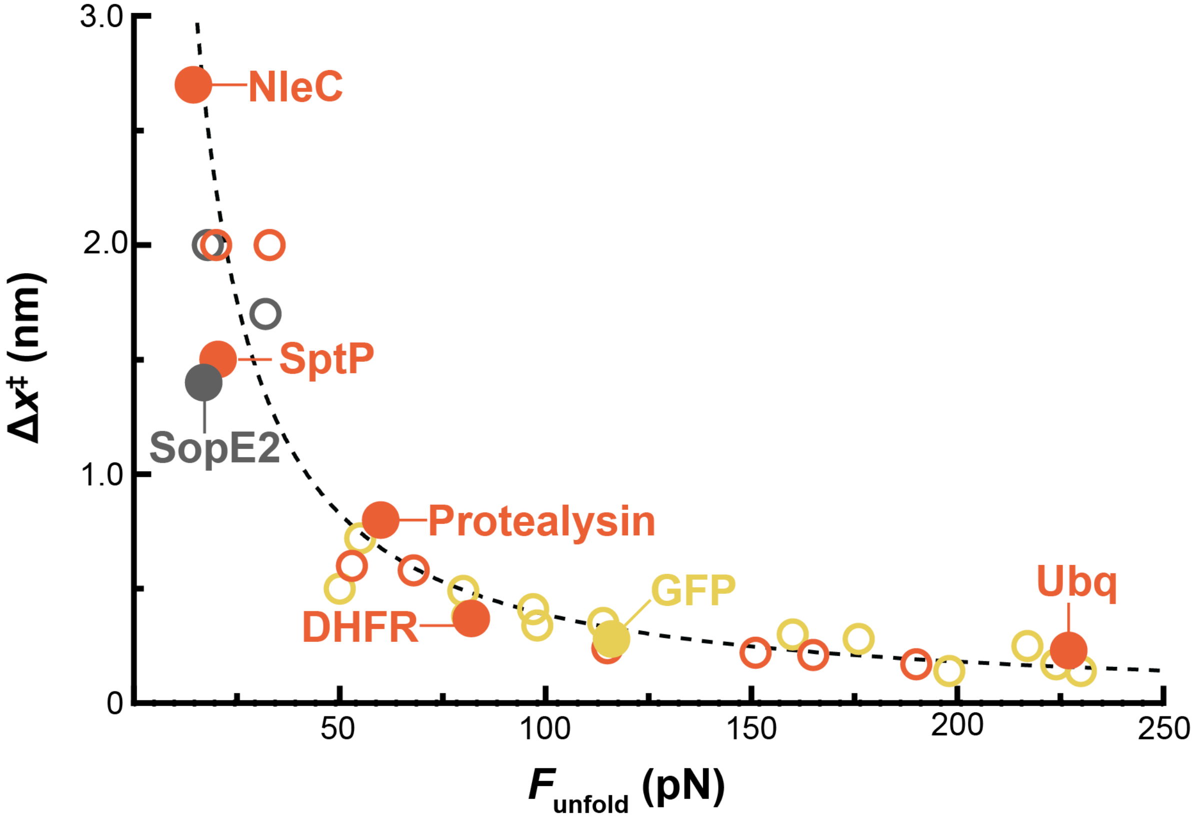


**Figure S3.** Mechanical properties of effectors compared to non-effector homolog and characterized proteins. The distance to the transition state, $\Delta x^{\ddagger}$, was plotted as a function of unfolding force ($F$) at a retraction velocity of 600 nm/s. Mechanical properties of reference proteins were compiled by Hoffman et al (8) and mechanical properties of SopE2 and SptP were obtained from reference (7). Data point shapes are color coded by their structural class: all α-helical (grey), αβ (orange), and all β-sheet (yellow). Effectors (NleC, SopE2 and SptP), non-effector homolog (protealysin), and proteins known to impair secretion (GFP, ubiquitin and DHFR) are highlighted with filled in markers. The overall hyperbolic relationship is well-described by fitting with the Bell-Evans model [dashed lines, (9)], $F_{unfold} = (\Delta x^{\ddagger}/k_B T)^{-1} ln[r \Delta x^{\ddagger}/(k_0 k_B T)]$, when the unfolding force is expressed in terms of distance to the transition state, while the loading rate ($r$ = 200 nm/s) and zero-force unfolding rate ($k_0$ = 0.2 $s^{-1}$) are held constant.

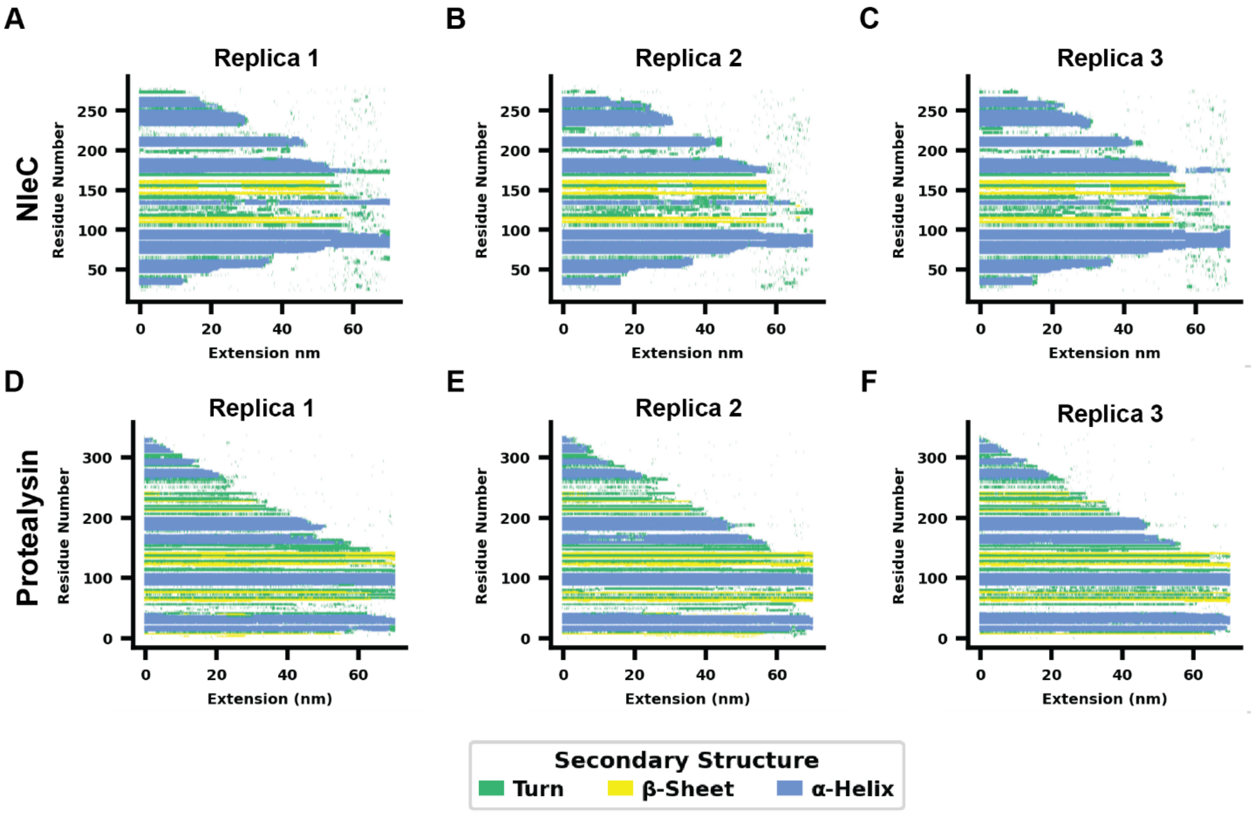


**Figure S4.** Secondary structure evolution plot for NleC and protealysin SMD simulations. Secondary structure elements (turn: green; β-sheet: yellow; α-helix: blue) monitored over simulation replicates. (A–C) NleC 350 ns simulation replicates show a reproducible unfolding pathway that initiates at the N-terminus followed by unfolding of both the N- and C-terminus. (D–F) Protealysin 250 ns simulation replicates depict a consistent unfolding pathway that initiates at the C-terminus.

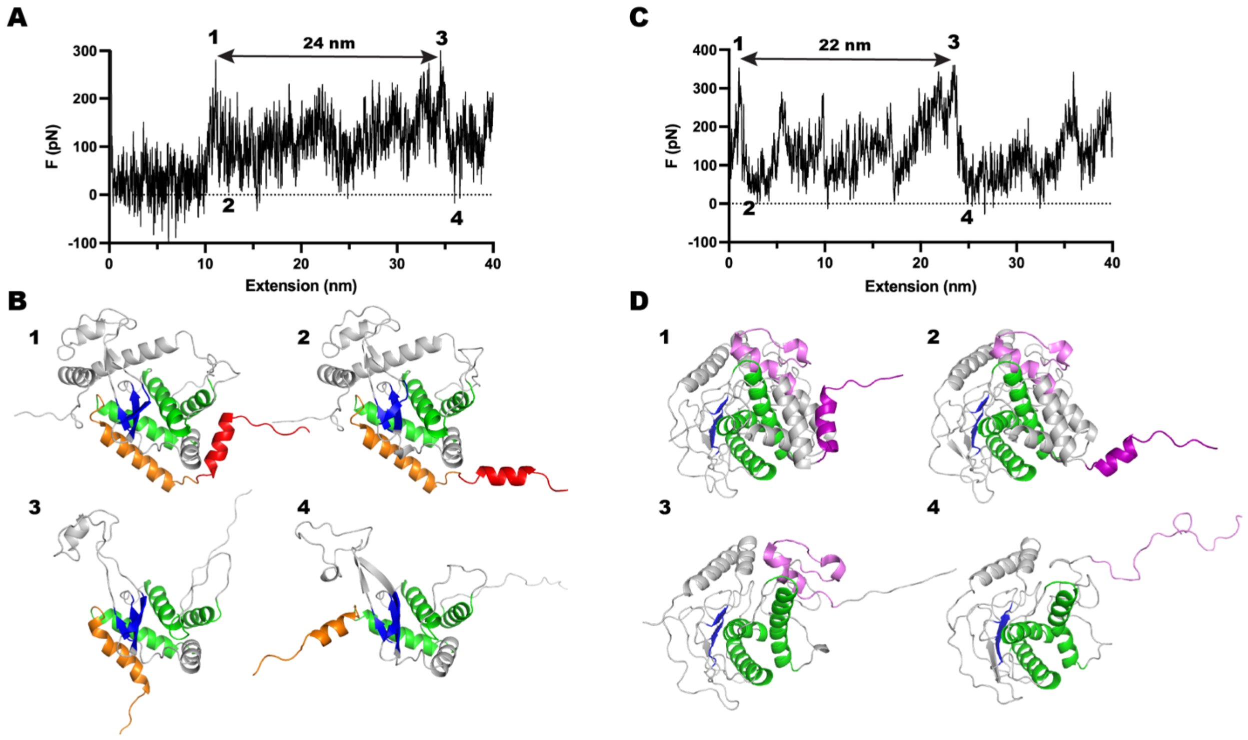


**Figure S5.** Force-induced unfolding behavior of NleC and protealysin in SMD simulations. (A) Representative force extension curve of a 350 ns SMD simulation replicate of NleC unfolding. The major force peaks (1 and 3) are separated by 24 nm. The force minima following the major peaks are indicated (2 and 4). (B) SMD simulation frames depict NleC unfolding corresponding to regions 1-4 indicated in (A). NleC unfolding initiates by release of the most N-terminal helix (red; 1→ 2). Prior to the second major force peak (2→3) at 34 nm, the c-terminus unfolds (grey). The second major force peak corresponds to the release of the second N-terminal helix (orange; 3→4). (C) Representative force extension curve of a 250 ns SMD simulation replicate of protealysin unfolding. The major force peaks (1 and 3) are separated by 22 nm. The force minima following the major peaks are indicated (2 and 4). (D) Protealysin unfolding initiates with the release of the C-terminal helix (pink; 1→2). Prior to the second major force peak (2→3) at 24 nm, the C-terminus continues to unfold (grey). The second major force peak corresponds to the unfolding of a helix-turn-helix motif (pink; 3→4).

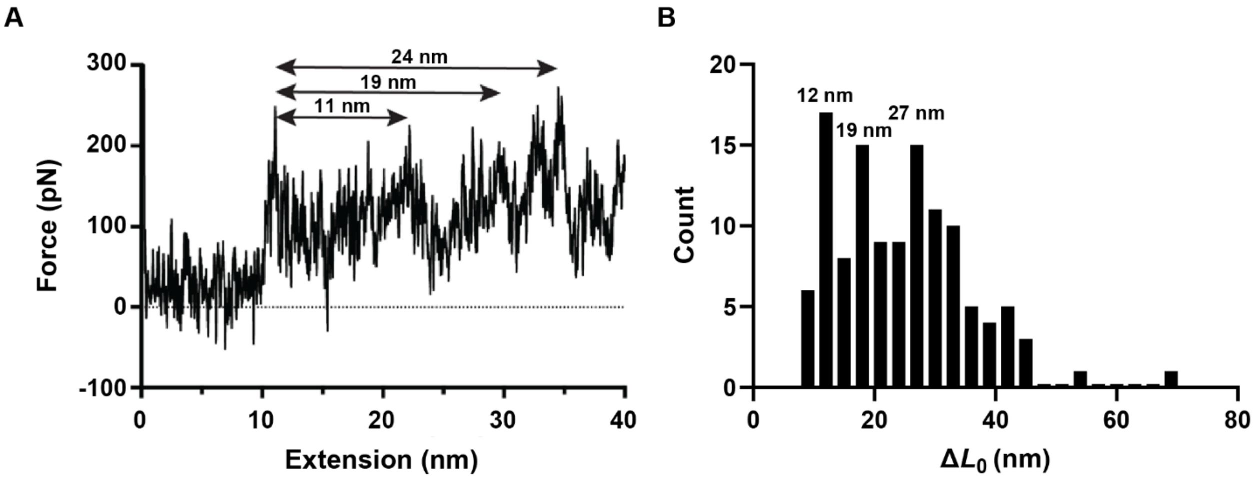


**Figure S6.** Unfolding transitions of NleC in SMD simulations and AFM assays. Representative SMD simulation FEC highlights the change in extension between the first major unfolding peak to minor (11 and 19 nm) and major force peaks (24 nm). (B) Distribution of first $\Delta L_0$ for all NleC unfolding events in AFM assays that proceed through one intermediate. The most abundant $\Delta L_0$s observed, 12, 19 and 27 nm, are indicated Histogram generated from N = 129 unfolding events at *v* = 100, 400, 800, 1600 and 3200 nm/s.

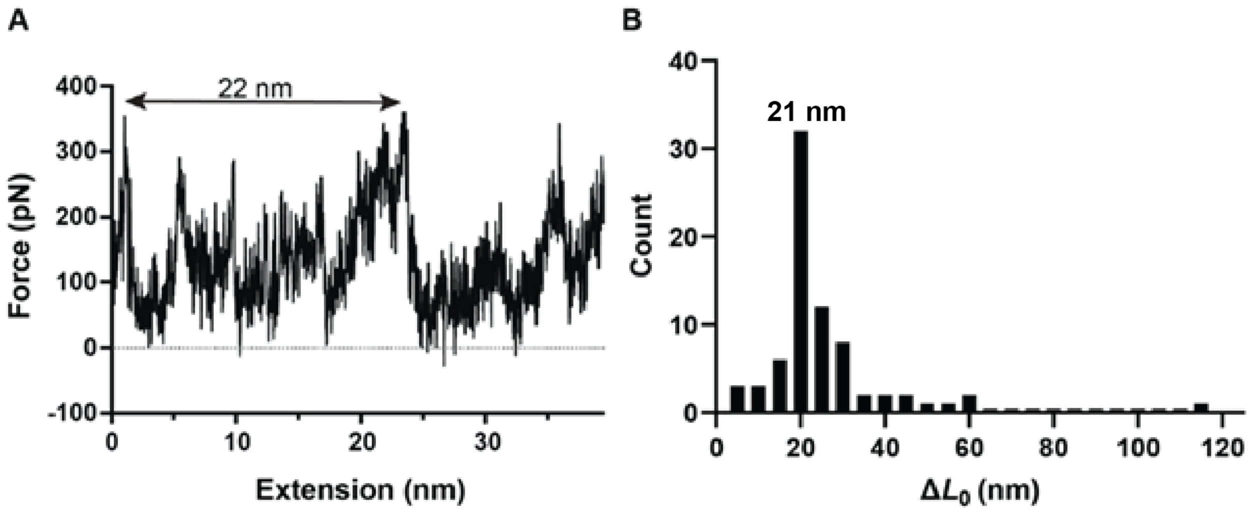


**Figure S7.** Unfolding transitions of protealysin in SMD simulations and AFM assays. (A) Representative SMD simulation FEC highlights the change of extension between the first and second major unfolding peaks (22 nm) (B) Distribution of first change in contour length for all protealysin unfolding events that proceed through one intermediate in AFM assays. The most abundant $\Delta L_0$ observed is 21 nm. Histogram generated from N = 75 unfolding events at $v$ = 30, 100, 300, 1000 and 3000 nm/s.

**Table S1.** Values from Bell-Evans model (3) fits of dynamic force spectra. Uncertainties are represented by the fit error.

| Protein | $\Delta x^{\ddagger}$ (nm) | $k_0$ ($s^{-1}$) |
|---|---|---|
| **NleC** | 2.7 ± 0.3 | $0.9 \pm 1.0 \times 10^{-2}$ |
| **Protealysin** | 0.7 ± 0.1 | $4.7 \pm 3.5 \times 10^{-3}$ |

**Table S2.** PCR Primers

| Primer | Sequence (5'→3') | Purpose |
|---|---|---|
| U2418 | CTCCGGGCTCCGGTTCTGGTATTGCTCCTAATCGTGCTG | NleC into pMS-1480 for Gibson assembly |
| L2418 | TTTGGTACCACAGAGCCGGAATACTCTGTGAAATCAGGAC | NleC into pMS-1480 for Gibson assembly |
| U2515 | ACCTTCACCGTTACCGAAGGCTCCGGTTCTGGTCCGACC | Protealyin into pMS-1711 for Gibson assembly |
| L2483 | GTTTGGTACCACAGAGCCGGACGCCACCCCTACCTGATGCCA | Protealyin into pMS-1711 for Gibson assembly |
| U2414 | ACCAGAACCGGAGCCCGG | Linearize pMS 1480 for Gibson assembly |
| L2414 | TCCGGCTCTGTGGTACCAAAC | Linearize pMS 1480 for Gibson assembly |
| U2442 | GGTATTGCTCCTAATCGTGCTGAAAATGCC | NleC into pMS-984 – Adds Gly |
| L2438 | TCGACTAGGAATACTCTGTGAAATCAGGACTCGCCTC | NleC into pMS-984 – Adds stop codon |

#### References


1. M. Schlierf, F. Berkemeier, M. Rief, Direct observation of active protein folding using lock-in force spectroscopy. *Biophys J* **93**, 3989-3998 (2007).
2. Y. Cao, H. Li, Polyprotein of GB1 is an ideal artificial elastomeric protein. *Nat Mater* **6**, 109-114 (2007).
3. M. S. Bull, R. M. Sullan, H. Li, T. T. Perkins, Improved single molecule force spectroscopy using micromachined cantilevers. *ACS Nano* **8**, 4984-4995 (2014).
4. J. K. Faulk, D. T. Edwards, M. S. Bull, T. T. Perkins, Improved Force Spectroscopy Using Focused-Ion-Beam-Modified Cantilevers. *Methods Enzymol* **582**, 321-351 (2017).
5. R. Walder *et al.*, Rapid Characterization of a Mechanically Labile alpha-Helical Protein Enabled by Efficient Site-Specific Bioconjugation. *J Am Chem Soc* **139**, 9867-9875 (2017).
6. J. Yin, A. J. Lin, D. E. Golan, C. T. Walsh, Site-specific protein labeling by Sfp phosphopantetheinyl transferase. *Nat Protoc* **1**, 280-285 (2006).
7. M. A. LeBlanc, M. R. Fink, T. T. Perkins, M. C. Sousa, Type III secretion system effector proteins are mechanically labile. *Proc Natl Acad Sci U S A* **118** (2021).
8. T. Hoffmann, K. M. Tych, M. L. Hughes, D. J. Brockwell, L. Dougan, Towards design principles for determining the mechanical stability of proteins. *Phys Chem Chem Phys* **15**, 15767-15780 (2013).
9. E. Evans, K. Ritchie, Dynamic strength of molecular adhesion bonds. *Biophys. J.* **72**, 1541-1555 (1997).
